# Genome sequencing of ion-beam-induced mutants facilitates detection of candidate genes responsible for phenotypes of mutants in rice

**DOI:** 10.1101/666677

**Authors:** Yutaka Oono, Hiroyuki Ichida, Ryouhei Morita, Shigeki Nozawa, Katsuya Satoh, Akemi Shimizu, Tomoko Abe, Hiroshi Kato, Yoshihiro Hase

**Author notes:** Corresponding Author: Yutaka Oono 1233 Watanuki, Takasaki, 370-1292 JAPAN (YO).

## Abstract

Ion beams are physical mutagens used for plant and microbe breeding that cause mutations via a distinct mechanism from those of chemical mutagens or gamma rays. We utilized whole-exome sequencing of rice DNA in order to understand the properties of ion beam-induced mutations in a genome-wide manner. DNA libraries were constructed from selected carbon-ion-beam-induced rice mutants by capturing with a custom probes covering 66.3 M bases of nearly all exons and miRNAs predicted in the genome. A total of 56 mutations, including 24 single nucleotide variations, 23 deletions, and 5 insertions, were detected in five mutant rice lines (two dwarf and three early-heading-date mutants). The mutations were distributed among all 12 chromosomes, and the average mutation frequency in the M1 generation was estimated to be 2.7 × 10^-7^ per base. Many single base insertions and deletions were associated with homopolymeric repeats, whereas larger deletions up to seven base pairs were observed at polynucleotide repeats in the DNA sequences of the mutation sites. Of the 56 mutations, six were classified as high-impact mutations that caused a frame shift or loss of exons. A gene that was functionally related to the phenotype of the mutant was disrupted by a high-impact mutation in four of the five lines tested, suggesting that whole-exome sequencing of ion-beam-irradiated mutants could facilitate the detection of candidate genes responsible for the mutant phenotypes.

## 1. Introduction

Ion beams are charged particles produced by particle accelerators that use electromagnetic fields. As with other ionizing radiations, ion beams cause damage to DNA molecules in living organisms and have been used as physical mutagens in plant and microbe breeding [1–4]. Ion beams are characterized by the deposition of a high energy transfer per unit length (linear energy transfer, LET) and are believed to induce mutations as a consequence of distinct biological effects from low LET radiation such as gamma-rays and electrons. Indeed, during the screening of mutants from irradiated explants of carnations, ion beams induced a wider variety of mutants with respect to flower color and shape than gamma-rays and X-rays [5]. A qualitative difference in flower color variations in an ion-beam-irradiated population has also been observed in chrysanthemums [1]. In contrast, Yamaguchi et al. reported no remarkable differences in the mutation spectra between ion-beam and gamma-ray irradiation [6].

Characterization of ion beam-induced mutations in plant DNA was conducted using several approaches in Arabidopsis, a model plant for plant molecular genetics. The most common approach for characterizing germline mutations is the isolation of mutants deficient in well-characterized marker genes responsible for visible phenotypes such as seed color (*tt*) and leaf trichome morphology (*gl*), followed by analysis of the DNA sequence of the marker gene [7–10]. The isolation of mutants or tissue sectors of well-characterized marker genes is also useful for characterizing ion-beam-induced somaclonal mutations by detecting the loss of heterozygosity [11–13]. An alternative method that can eliminate the isolation of mutant plants is the use of the *Escherichia coli* ribosomal protein small subunit S12 (*rpsL*) mutation detection system [14, 15]. In this system, plants containing the *rpsL* transgene are irradiated, and genomic DNA from irradiated transgenic plants are introduced into *E. coli*. Mutated nonfunctional *rpsL* DNA fragments can be recovered from drug-resistant *E. coli* colonies. Previous experiments using these methods suggest that 1) ion beams induce various types of mutations into plant DNA, such as base substitutions, DNA insertions and deletions (InDels), inversions (INVs), and chromosomal translocations [8, 10]; 2) ion beams induce germline mutations with a higher ratio of rearrangements such as InDels greater than 100 bp, inversions, translocations, and total deletions of the marker gene than electron beams, which induce mostly point-like mutations such as base substitutions and InDels less than100 bp [8]; 3) the size of the deletion induced by ion beams positively correlates with the degree of LET [12,16,17]; and 4) most mutants obtained with a pollen-irradiation method have large deletions > 6 Mb, most of which are not transmittable to the next generation [11].

Recent advances in genome analysis technologies have allowed for more quantitative evaluation of mutations at the genomic level without bias arising from genomic positioning and of the functional significance of DNA sequences of a specific marker gene. A mutation-accumulating experiment combined with whole genome analysis with next-generation sequencing revealed an estimated spontaneous mutation rate in Arabidopsis of 7 × 10^−9^ base substitutions per site per generation [18]. Whole genome sequencing analysis of ion beam-irradiated Arabidopsis DNA suggested that 200-Gy carbon ion beams increased the mutation rate nearly 47-fold compared to spontaneous mutations [19]. In addition, Arabidopsis whole genome sequencing has demonstrated that the frequency and type of mutations induced by ion beams are affected by the LET of ion beams [20] and the physiological status of irradiated tissues [21].

Rice is one of the most important crops for humans and is a major target for ion-beam breeding strategies [22–26]. Several genes that cause mutant phenotypes were successfully cloned by map-based cloning from ion-beam-induced rice mutants, and the mutation sites in the genes were characterized [23, 26]. However, due to a relatively large genome size compared to Arabidopsis, the characterization of ion beam-induced mutations at the genomic level was not achieved in rice until very recently [27, 28]. In the present work, we used a whole-exome sequencing procedure to analyze the properties of induced mutations in selected rice mutants generated with carbon ion beams accelerated using the Takasaki Ion Accelerators for Advanced Radiation Application (TIARA) azimuthally varying field (AVF) cyclotron at TARRI, QST. We demonstrated that a relatively small number of induced mutations with a potentially high impact on protein function quickly narrowed down the candidate genes responsible for the mutant phenotypes.

## 2. Materials and Methods

### 2.1 Plant materials and ion beam irradiation

Rice seeds (*Oryza sativa* L. cv Nipponbare, NPB) used in this work were harvested in RBD, NICS, NARO (Hitachi-ohmiya, Japan). Water content in the rice seeds were adjusted to 12–13% by keeping the rice seeds in a plant growth chamber (BIOTRON LPH200, NK system, Japan) at 24°C and 60% RH for 4 d just before irradiation. Then, the seeds were irradiated with 10–150 Gy (at 10 Gy intervals) of 25.9 MeV/u ^12^C^6+^ ions (LET on surface: 76 keV/μm) accelerated by an AVF cyclotron at TIARA, TARRI, QST (Takasaki, Japan). The irradiated seeds were sown on soil to produce M1 plants in a greenhouse at TARRI, QST. The dose response for the survival rate and plant height were measured one month after sowing. Plants that produced green leaves were considered survivors and length from the ground to the tip of the leaf blade at the highest position in stretched tillers was measured as plant height. The seeds treated with 40 Gy of ^12^C^6+^ ions were used for a genetic mutant screen. M2 or M3 seeds were harvested from every individual mature M1 or M2 plant, respectively. For selection of mutant candidates, M2 and M3 plants were grown in the greenhouse at TARRI, QST or in an experimental paddy field at RBD, NICS, NARO (36°52′45″N 140°39′81″E). Some mutant lines were backcrossed to un-mutagenized NPB. The resulting seeds (F1) were germinated and grown in the greenhouse to generate F2 seeds, which were used for genetic linkage analysis between the mutations and the mutant phenotype.

### 2.2 DNA extraction, whole exome capturing, and sequencing

Genomic DNA for whole exome capture was extracted from rice leaves of single individual plants using MagExtractor –Plant Genome– (Toyobo, Japan) according to the manufacturer protocol. One microgram of genomic DNA from individual samples was fragmented using a Covaris S220 instrument (Covaris Inc, USA) and resulting DNA fragments were purified with AMPure XP reagent (Beckman Coulter, USA). Barcoded sequencing libraries were prepared using NextFlex Rapid DNA-Seq Kits (Bioo Scientific, USA) according to the manufacturer’s protocol. After double-sided size-selection with AMPure XP, pre-capture amplification was performed for 12 cycles with the supplied primers 1 and 2. The Os-Nipponbare-Reference-IRGSP-1.0 sequences and annotations were used as a reference (https://rapdb.dna.affrc.go.jp/download/irgsp1.html; as of 2015/3/31). A custom whole-exome-capturing probe library was designed and synthesized as a SeqCap EZ Developer Library (Roche Diagnostics, USA). This library covered 66.3 Mb of 107,616 rice genome regions that consisted of all predicted exons in the IRGSP-1.0 annotations, including 100 bp each of the 5’- and 3’-franking regions, as well as known miRNA regions including 700 bp and 300 bp of the upstream and the downstream regions, respectively. After 16–20 h of hybridization at 47°C, the hybridized DNA fragments were captured with streptavidin-coupled magnetic beads (Dynabeads M-280 streptavidin, Thermo Fisher Scientific, USA), washed, and amplified by PCR using KAPA HiFi Hotstart Ready Mix (KAPA Biosystems) with the same primers described above. The whole exon-enriched libraries were sequenced on a HiSeq 4000 instrument (Illumina, USA) in PE100 mode.

Data analysis was performed with the bioinformatics pipeline described previously [27]. Briefly, sequencing reads were mapped to the reference sequences using the BWA-MEM program version 0.7.15 (http://bio-bwa.sourceforge.net/) with default parameters, followed by manipulation with SAMtools version 1.3.1 (http://samtools.sourceforge.net/), Picard software package version 2.3.0 (https://broadinstitute.github.io/picard/) was used to sort the reads and mark and remove PCR duplicates. GATK software package version 3.7.0 (https://software.broadinstitute.org/gatk/) was used for local realignment and base quality recalibration. Variant calling was carried out with a combination of the UnifiedGenotyper tool in GATK, Pindel version 0.2.5a8 (http://gmt.genome.wustl.edu/packages/pindel/), and Bedtools version 2.26.0 (https://github.com/arq5x/bedtools2/) programs. To remove intra-cultivar polymorphisms between the reference sequences and our parental NPB line used for the irradiation experiments, the raw variant calling results among the mutants were compared and the variants present in only one mutant line were extracted as the line-specific (i.e. real) mutations. For the GATK and Pindel results, the detected line-specific mutations were further filtered out using the following criteria: 1) a mutation that was supported with less than 10 reads was removed to eliminate false-positives due to an insufficient number of mapped reads; and 2) a homozygous mutation was removed if reads with the same mutation accounted for more than 5% of the total reads found in equal or more than half of the homozygous lines for the other allele (i.e. unmutated allele). Mutations detected with each program were merged together by the normalized chromosomal positions (start and end coordinates), and duplicated mutation records that were detected by multiple programs were removed from the final result. In the Bedtool analysis, regions covering less than 20% in one individual line but 100% in all the other lines were extracted. To increase reliability, regions with less than an average of 10 reads in all lines with a wild-type homozygous allele were excluded. The predicted effect and location of the mutations were annotated using the SnpEff program version 4.1b (http://snpeff.sourceforge.net/). After variant calling against the whole genome sequence, only mutations located within the exome target regions of whole-exome capturing were extracted and outputted in variant calling format (VCF). The identified mutations were checked visually using the integrative genomics viewer (IGV) (http://www.broadinstitute.org/igv/).

To confirm the mutation by Sanger sequencing, the DNA fragment of the mutation site was amplified by PCR using Takara Ex Taq polymerase (Takara Bio, Japan) followed by the dye terminator cycle sequencing reaction with the BigDye Terminator v3.1 Cycle Sequencing Kit (Thermo Fisher Scientific). The resulting samples were analyzed with a 3500 Genetic Analyzer (Thermo Fisher Scientific). The primers used for PCR and Sanger sequencing are listed in S1 Table.

### 2.3 Estimation of the genome-wide mutation frequency

Mutation frequencies (MFs) were calculated by dividing the total number of mutations detected in individual plant DNA by the number of bases that achieved 10X or greater coverage in the captured target region (Table1). Assuming all mutations arise as heterozygotes in the M1 generation and are inherited according to Mendelian characteristics, all homozygous mutations in M2 and later generations were transmitted to the next generation, but a quarter of the heterozygous mutations were not. Therefore, the number of mutations in M1 plants (N1) was estimated as

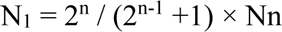

where n is the generation and Nn is the number of mutations in the Mn diploid genome.

## 3. Results and Discussion

### 3.1 Preparation of an ion-beam irradiated rice population and mutant selection

We chose the rice cultivar Nipponbare (NPB) in this work because a high-quality whole genome sequence is available [30]. Since the water content of a seed affects its radiation sensitivity [31], we adjusted the water contents of the seeds to 12 – 13% before irradiation. To estimate the biological effectiveness of the carbon ion beam on the NPB seeds, a dose response for the survival rate and plant height were determined (Fig. 1A). A shoulder or slightly lower dose in the survival curve was empirically considered to be optimal for obtaining a large number of mutants [1,6,22] and therefore, the absorbed dose chosen for the genetic screen in this work was 40 Gy, which was approximately 2/3 of the shoulder dose (approximately 60 Gy) for the survival curve and slightly over the shoulder dose for the dose response curve of plant height.

**Fig. 1.**
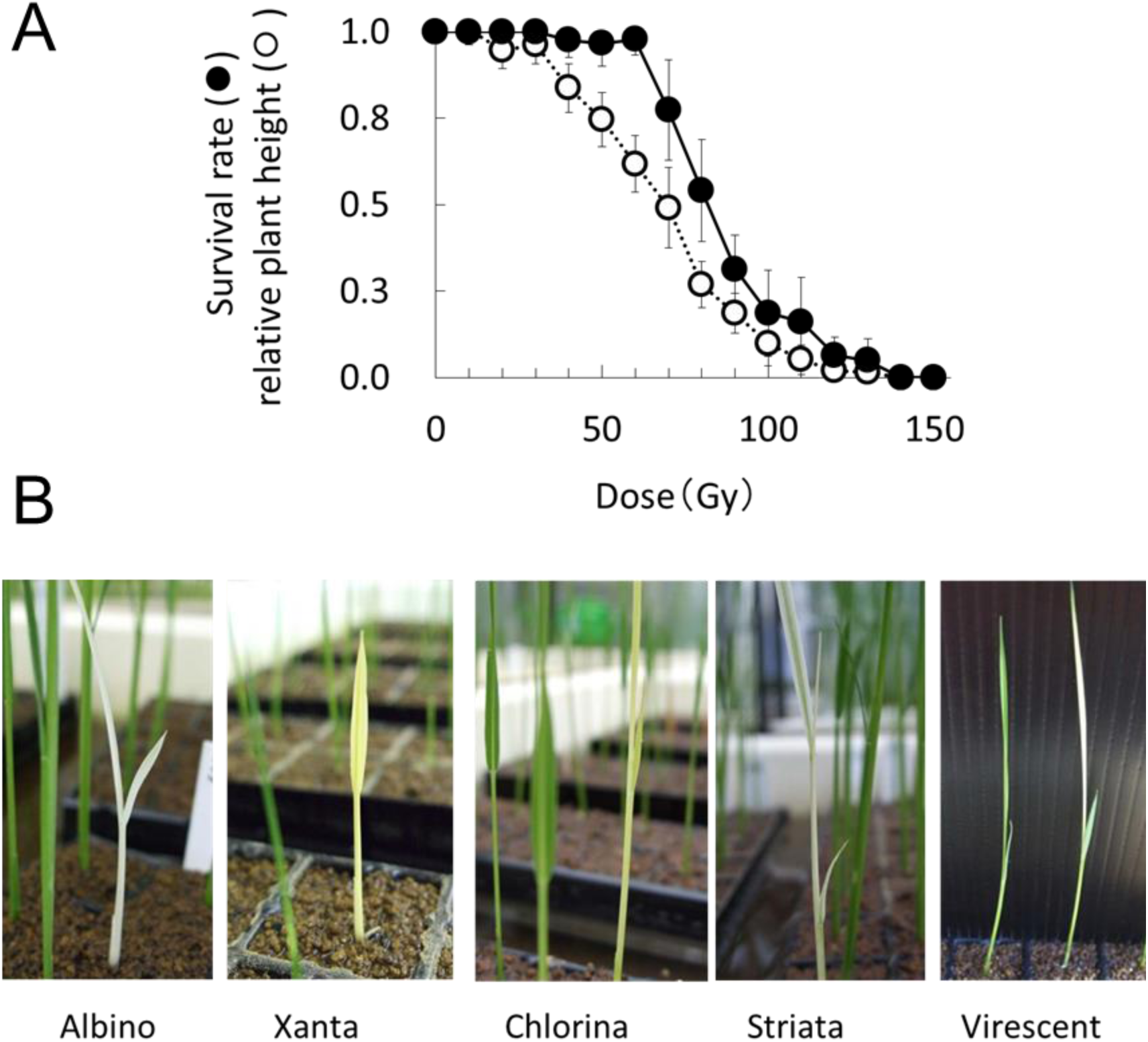
Estimating the effect of carbon ion beams on rice. (A) Dose response of the survival rate and relative plant height of NPB rice seeds irradiated by ^12^C^6+^ ion beams. Plotted values represent as mean values against unirradiated controls (0 Gy). Error bars indicate standard error (SE). Data are from five replicates with 10 seedlings per treatment. (B) Chlorophyll-related mutant phenotypes. The number of mutants in the M2 population was counted to estimate the mutagenic effect of the 40 Gy carbon ion beam irradiation treatment.

To estimate the mutagenic effect of 40 Gy irradiation treatment, M2 seeds (4 – 10 seeds per M1 line) from 2,039 individual M1 plants were sown on soil and chlorophyll mutants were identified at the seedling stage. In this pilot study, several types of chlorophyll mutants with a pale or white leaf color were identified (Fig 1B). The appearance rate of the chlorophyll mutants was 6.6% (134 mutants in 2,039 M1 lines), suggesting that the carbon-ion-beam irradiation effectively induced mutations in the rice genome as expected. We also screened other types of mutants with altered phenotypes from this M2 population in the green house and paddy field. In this work, we focused on two dwarf (lines “3098” and “885”) and three early-heading-date (lines “786-5”, “IRB3517-3”, and “IRB3790-2”) mutants (Fig. 2) that had the same phenotypes in the next (M3) generation. The mutant lines “3098” and “885” were identified in the greenhouse, whereas lines “IRB3517-3”, and “IRB3790-2” were found in the paddy field at RBD (formerly the Institute of Radiation Breeding (IRB)). Line “786-5” was initially found in the green house as a mutant candidate with wide leaves and later developed an early heading phenotype that was detected in the paddy field (Fig. 2C and D). Homozygosity of the mutant phenotypes was confirmed in the M3 generation of all the lines except “IRB3790-2”, for which we failed to obtain a homozygous population in the M3 and M4 generations (Fig. 2D). Leaf tissues excised from a single M3 plant from each line except “3098”, for which the leaf tissue was from a single M2 plant, were used for genomic DNA extraction and whole-exome sequencing analysis.

**Fig 2.**
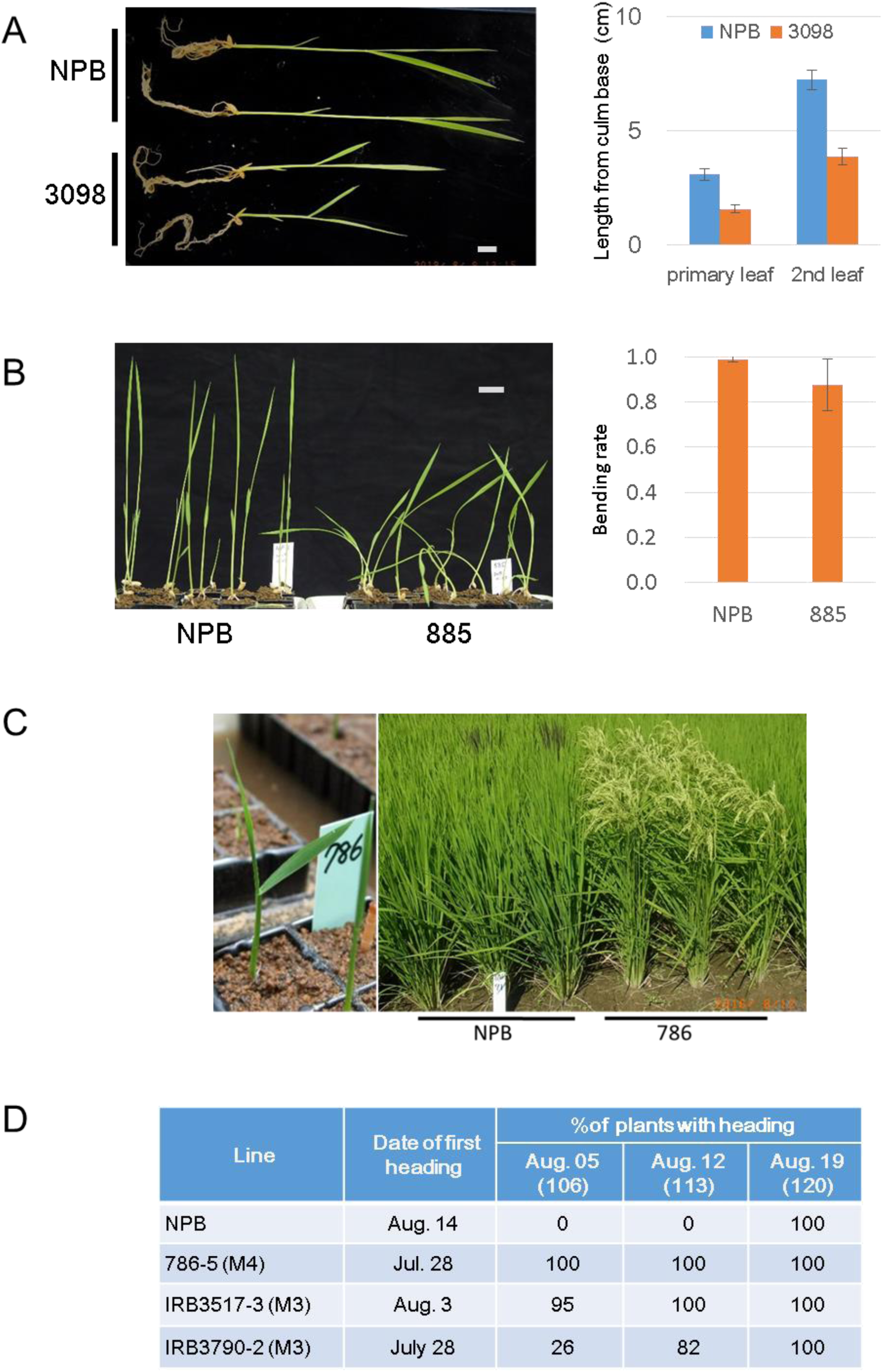
Dwarf and early-heading-date mutant lines identified from irradiated rice. (A) Six-teen-d-old NPB control and “3098” (M4) plants grown in the green house. The average distance from culm base are displayed in the bar chart. The values of the average distances from culm base to the lamina joint of primary and secondary leaves were 3.1 ± 0.2 and 7.3 ± 0.4 cm for NPB and 1.6 ± 0.2 and 3.9 ± 0.4 cm for “3098”, respectively. The values were significantly different between NPB and “3098” for both measurements (P < 0.01 in *t*-test, n = 8). Scale bar = 1 cm. (B) Ten-d-old plants of NPB control and “885” (M5) grown in a plant growth chamber (12 h light/12 h dark light condition). The bending rate was calculated by dividing the linear distance from culm base to leaf tip by the stretched length from culm base to leaf tip of the same plant (0.99 ± 0.01 for NPB and 0.88 ± 0.11 for “885”, P < 0.05 in *t*-test, n = 8). Scale bar = 1 cm. (C) Left panel: The wide leaf phenotype of a “786” seedling grown in the green house. Right panel: NPB control and “786” (M4) rice growing in the paddy field. One-month-old seedlings were transferred to the paddy field on May 20, 2016. The photograph was obtained on August 12, 2016, when all “786” but not the NPB rice had completed heading initiation. (D) Heading dates of NPB, “786-5”, “IRB3517-3”, and “IRB3790-2” were recorded in the paddy field in 2016. One-month-old seedlings were transferred to the paddy field on May 20, 2016. The date of first heading in the population and the ratios of plants with heading recorded on August 05, 12, and 19 are shown. Numbers in parenthesis indicate days after sowing. The M4, M3, and M3 populations were used for “786-5”, “IRB3517-3”, and “IRB3790-2”, respectively.

### 3.2 Whole-exome captured sequencing in rice

We subjected all five mutants to whole-exome capture and massive parallel sequencing analysis to comprehensively identify mutations introduced by the carbon-ion beam irradiation. Whole-exome capture was followed by Illumina sequencing and we obtained the sequence data for 61.0 – 69.9 million unique reads with 6.08 – 6.96 billion unique bases aligned to the reference genome (Table 1). The number of bases mapped on the target region was 2.38 – 2.78 billion. The mean target coverage was 36 – 42, and the percentage of all target bases with 10 times or greater coverage was 95.4 – 96.0% (Table 1). Using the pipeline described in Materials and Methods, GATK, Pindel, and Bedtools programs initially identified 61 line-specific mutations in the target regions. By manual verification of VCF files, two or three mutations that were overlapped or located adjacent to one another were considered to be one mutation and categorized as a replacement (RPL) (mutations #12, 13, 37 in Table 2, S2 Table and Fig. 3). In the Bedtools program, one large deletion was identified on chromosome 3 of the “IRB3517-3” genome. This large deletion was also identified by the Pindel program (mutation # 38) and consequently, the number of mutations was reduced to 56 (Tables 2 and 3). To verify the accuracy of the mutation calling in the pipeline, DNA fragments corresponding to 25 selected mutations (Table 2), including the junction sites of a large deletion (#38) and inversion (#52), were amplified by PCR and reanalyzed by Sanger sequencing. The sequences of all the selected mutations from the pipeline were confirmed except for the 3-bp shift at one of the junction sites of the #52 inversion, suggesting that the results of our exome analysis were reliable, with respect to false positives.

**Fig. 3.**
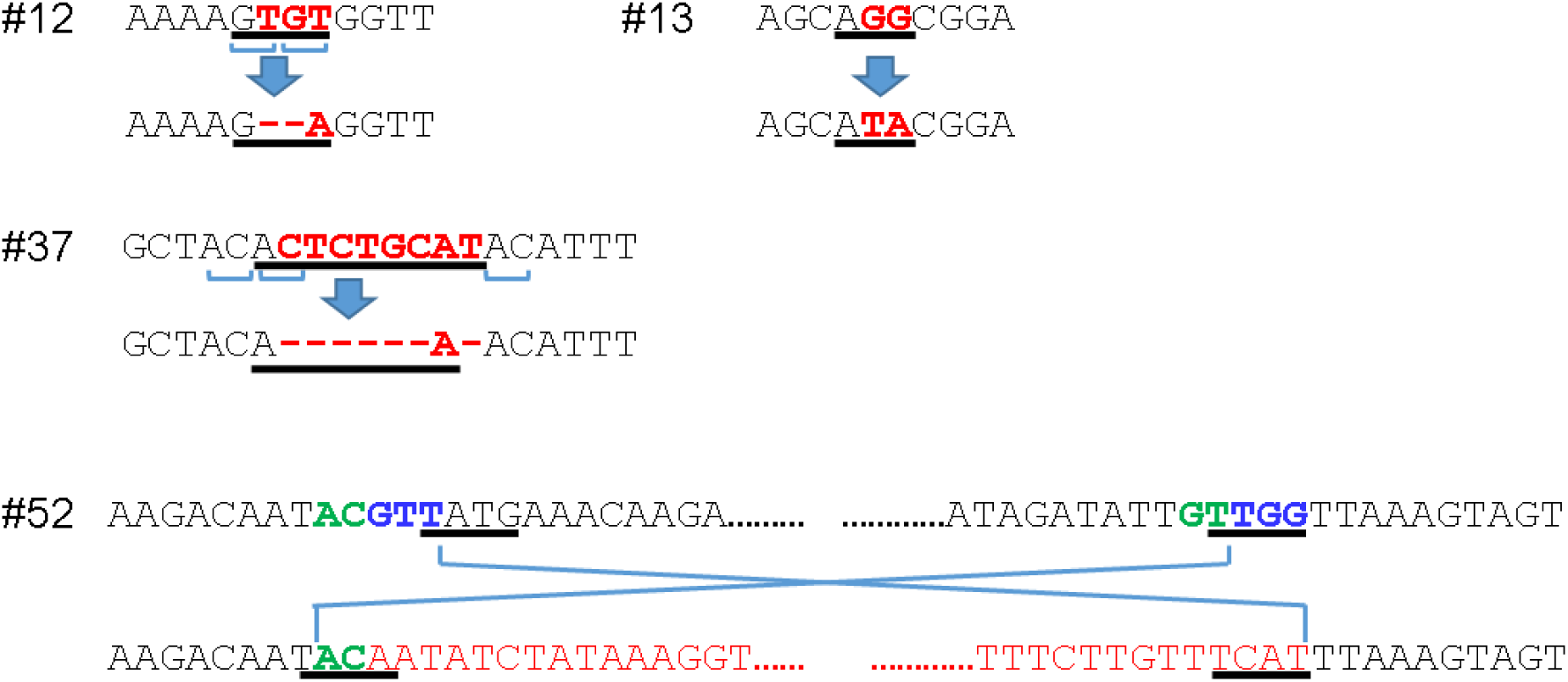
Nucleotide sequences in the mutation site of three RPLs and junction sites of the 535-kb INV. Sequences in the NPB control (top) and mutant (bottom) rice samples are shown with underlined regions indicating the nucleotides of “Reference” and “Alternatives” from S2 Table. For mutations #12 and #37, altered nucleotides between the NPB control and mutants are shown in red. The dinucleotide repeats in control DNA at #12 and #37 mutation sites are indicated by top-opened brackets. For mutation #52, green and blue letters indicate overlapped and deleted nucleotides in junction sites, respectively. Inverted sequences, excluding the overlapped nucleotides, are shown in red.

**Table 1.** Summary of the whole-exome captured sequencing

|  | Line |  |  |  |  | Mean | SD |
| --- | --- | --- | --- | --- | --- | --- | --- |
|  | 3098 | 786 | 885 | IRB3517-3 | IRB3790-2 |  |  |
| Reference genome size (bases) | 373,245,519 | 373,245,519 | 373,245,519 | 373,245,519 | 373,245,519 |  |  |
| Size of target (bases) | 66,258,908 | 66,258,908 | 66,258,908 | 66,258,908 | 66,258,908 |  |  |
| No. of reads aligned | 65,509,961 | 61,048,713 | 69,002,757 | 69,858,886 | 67,703,204 | 66,624,704 | 3,521,809 |
| No. of bases aligned | 6,527,192,732 | 6,082,274,964 | 6,874,937,097 | 6,959,746,867 | 6,745,199,762 | 6,637,870,284 | 350,806,394 |
| No. of bases mapped<br>on targeted region | 2,684,758,131 | 2,384,714,814 | 2,637,344,060 | 2,776,001,880 | 2,734,422,333 | 2,643,448,244 | 153,731,509 |
| Mean target coverage | 40.52 | 35.99 | 39.80 | 41.90 | 41.27 | 39.90 | 2.32 |
| No. of bases 10X <sup>1</sup> | 63,362,069 | 63,220,208 | 63,574,097 | 63,611,931 | 63,417,925 | 63,437,246 | 159,979 |
| Fraction target bases 10X <sup>2</sup> | 0.956 | 0.954 | 0.959 | 0.960 | 0.957 | 0.957 | 0.002 |

214 <sup>1</sup> Number of bases with 10X or greater coverage in the target region.215 <sup>2</sup> The fraction of all target bases with 10X or greater coverage.

**Table 2.** List of mutations^1^

| Mutation No. | Line | CHR No. | Type after evaluation<sup>2</sup> | Annotation | Genotype <sup>3</sup> | Verification <sup>4</sup> |
| --- | --- | --- | --- | --- | --- | --- |
| 1 | 3098 | chr02 | INS (1) |  | 0\|1 |  |
| 2 | 3098 | chr02 | DEL (1) |  | 0\|1 |  |
| 3 | 3098 | chr03 | SNV | APG6/CLPB-P/CLPB3 | 0\|1 |  |
| 4 | 3098 | chr03 | SNV | DCL2A | 0\|1 |  |
| 5 | 3098 | chr04 | INS (7) |  | 1\|1 | ✓ |
| 6 | 3098 | chr05 | SNV | Diacylglycerol acylCoA acyltransferase | 0\|1 |  |
| **7** | **3098** | **chr05** | **DEL (128)** | **Guanine nucleotide-binding protein alpha-1 subunit** | **0\|1** | **✓** |
| 8 | 3098 | chr08 | DEL (1) |  | 0\|1 |  |
| 9 | 3098 | chr08 | INS (1) |  | 0\|1 |  |
| 10 | 3098 | chr12 | DEL (1) |  | 0\|1 |  |
| 11 | 786 | chr01 | SNV | Glutelin family protein | 0\|1 |  |
| 12 <sup>5</sup> | 786 | chr01 | RPL |  | 1\|1 | ✓ |
| 13 <sup>5</sup> | 786 | chr03 | RPL | Predicted protein | 1\|1 | ✓ |
| 14 | 786 | chr03 | SNV |  | 1\|1 |  |
| **15** | **786** | **chr03** | **DEL (5)** | **Phytochrome B** | **1\|1** | **✓** |
| 16 | 786 | chr06 | SNV |  | 1\|1 |  |
| 17 | 786 | chr07 | SNV |  | 1\|1 |  |
| **18** | **786** | **chr10** | **DEL (1)** | **Expansin/Lol pI family protein** | **1\|1** | **✓** |
| **19** | **786** | **chr12** | **DEL (1)** | **Nucleotide-binding, alpha-beta plait domain containing protein** | **0\|1** |  |
| 20 | 885 | chr01 | DEL (3) |  | 1\|1 |  |
| 21 | 885 | chr01 | DEL (1) |  | 0\|1 |  |
| 22 | 885 | chr01 | SNV |  | 1\|1 | ✓ |
| 23 | 885 | chr01 | DEL (46) | Soluble inorganic pyrophosphatase | 1\|1 | ✓ |
| 24 | 885 | chr02 | DEL (1) | Casein kinase I isoform delta-like | 0\|1 |  |
| 25 | 885 | chr03 | SNV |  | 0\|1 |  |
| 26 | 885 | chr03 | DEL (1) |  | 1\|1 | ✓ |
| 27 | 885 | chr04 | SNV |  | 1\|1 | ✓ |
| 28 | 885 | chr05 | SNV | Nucleotide-binding, alpha-beta plait domain containing protein | 0\|1 |  |
| 29 | 885 | chr05 | INS (1) |  | 0\|1 |  |
| 30 | 885 | chr05 | SNV | DNA glycosylase/lyase 701 | 1\|1 | ✓ |
| 31 | 885 | chr08 | SNV |  | 1\|1 |  |
| 32 | 885 | chr08 | DEL (6) |  | 1\|1 | ✓ |
| 33 | 885 | chr08 | DEL (4) |  | 1\|1 | ✓ |
| 34 | 885 | chr08 | SNV | Hypothetical conserved gene | 1\|1 | ✓ |
| 35 | 885 | chr10 | DEL (6) | CYP71Z8 | 1\|1 | ✓ |
| 36 | IRB3517-3 | chr01 | SNV | Cell division protein ftsH | 0\|1 |  |
| 37 <sup>5</sup> | IRB3517-3 | chr01 | RPL |  | 1\|1 | ✓ |
| **38** | **IRB3517-3** | **chr03** | **DEL (33.6k)** | **5 ORFs are deleted<sup>6</sup>** | **1\|1** | **✓** |
| 39 | IRB3517-3 | chr04 | SNV |  | 1\|1 |  |
| 40 | IRB3517-3 | chr06 | DEL (1) |  | 0\|1 |  |
| 41 | IRB3517-3 | chr06 | DEL (6) | Heavy metal ATPase | 1\|1 | ✓ |
| 42 | IRB3517-3 | chr07 | DEL (8) |  | 1\|1 | ✓ |
| 43 | IRB3517-3 | chr08 | SNV | Helix-loop-helix DNA-binding domain containing protein | 0\|1 |  |
| 44 | IRB3517-3 | chr09 | SNV | Phosphate/phosphoenolpyruvate translocator | 1\|1 | ✓ |
| 45 | IRB3517-3 | chr09 | SNV |  | 0\|1 | ✓ |
| 46 | IRB3517-3 | chr11 | SNV | Hypothetical gene | 1\|1 |  |
| 47 | IRB3517-3 | chr11 | SNV | NB-ARC domain containing protein | 1\|1 | ✓ |
| 48 | IRB3517-3 | chr12 | DEL (6) | Heterochromatin protein | 0\|1 |  |
| 49 | IRB3790-2 | chr01 | DEL (1) |  | 0\|1 |  |
| 50 | IRB3790-2 | chr03 | INS (1) |  | 0\|1 |  |
| 51 | IRB3790-2 | chr04 | SNV |  | 1\|1 |  |
| **52** | **IRB3790-2** | **chr06** | **INV (535k)** | **HEADING DATE 1<sup>6</sup>** | **0\|1** | **✓** |
| 53 | IRB3790-2 | chr07 | SNV | Esterase precursor | 0\|1 |  |
| 54 | IRB3790-2 | chr08 | DEL (4) |  | 1\|1 | ✓ |
| 55 | IRB3790-2 | chr09 | DEL (2) |  | 1\|1 |  |
| 56 | IRB3790-2 | chr12 | SNV |  | 1\|1 | ✓ |

<sup>1</sup> Mutations with a putative high impact on protein function are indicated in bold.
<sup>2</sup> Type of mutations are single nucleotide variation (SNV), deletion (DEL), insertion (INS), inversion (INV), and replacement (RPL). The size of INSs and DELs are indicated in parentheses.
<sup>3</sup> "0|1" is heterozygous and "1|1" is homozygous.
<sup>4</sup> Mutations verified by Sanger sequencing are marked with a checkmark. Primer sequences are shown in S1 Table.
<sup>5</sup> Independent mutation calls unified and counted as one RPL mutation because they overlapped or located consecutively (Fig. 3).
<sup>6</sup> Evaluated manually.

**Table 3.** Summary of the number of mutations and MF.

| Line name | Phenotype | Generation | # of mutation |  |  |  | MF | Estimated MF<br>in M1 | # of<br>high impact<br>mutation | Type of<br>high impact<br>mutation |
| --- | --- | --- | --- | --- | --- | --- | --- | --- | --- | --- |
|  |  |  | Total | SNV | InDel | Others |  |  |  |  |
| 3098 | Dwarf<br>Small grains | M2 | 10 | 3 | 7 | 0 | 1.6E-07 | 2.1E-07 | 1 | 128-bp DEL |
| 786 | Early heading date<br>Wide leaves<br>Broken flag leaf | M3 | 9 | 4 | 3 | 2 | 1.4E-07 | 2.3E-07 | 3 | 1-bp DEL x2,<br>5-bp DEL |
| 885 | Dwarf<br>Culm bending | M3 | 16 | 7 | 9 | 0 | 2.5E-07 | 4.0E-07 | 0 |  |
| IRB3517-3 | Early heading date | M3 | 13 | 7 | 5 | 1 | 2.0E-07 | 3.3E-07 | 1 | 33.6-kb DEL |
| IRB3790-2 | Early heading date | M3 | 8 | 3 | 4 | 1 | 1.3E-07 | 2.0E-07 | 1 | 535-kb INV |
| Total |  |  | 56 | 24 | 28 | 4 |  |  | 6 |  |
| Average |  |  | 11.2 |  |  |  |  | 2.7E-07 | 1.2 |  |

<sup>1</sup>MFs were calculated by dividing the total number of mutations detected in individual plant DNA samples by the number of bases with 10X or greater coverage in the captured target region (Table 1).

The 56 mutations consisted of 24 single nucleotide variations (SNV), 23 deletions (DELs), five insertions (INSs), one inversion (INV), and three RPLs (Tables 2 and 3). The average number of mutations per line was 11.2 ± 3.3 for all five mutant lines and 11.5 ± 3.7 for the four M3 lines. The mutations were distributed among all 12 chromosomes (Fig 4). The 1-bp DEL (#40) in “IRB 3517-3” and the 535-kb large INV (#52) in “IRB 3790-2” were located at the same region on the upper arm of chromosome 6. The mutation frequencies (MFs) were calculated to be 1.3 × 10^-7^ to 2.5 × 10^-7^ per base, and the MFs in the M1 generation were estimated to be 2.0 × 10^-7^ to 4.0 × 10^-7^ per base and, on average, 2.7 × 10^-7^ per base (Table 3).

**Fig 4.**
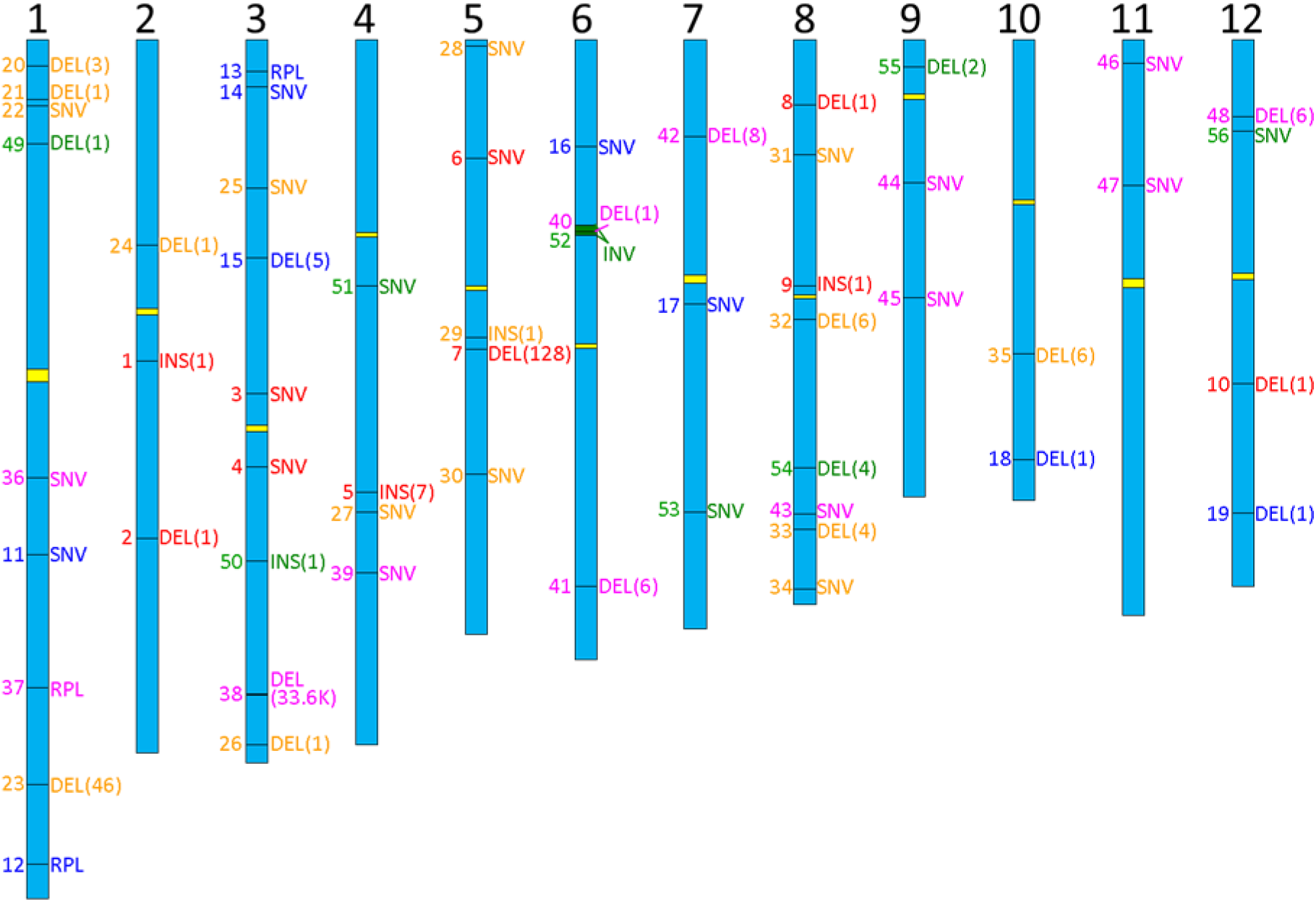
Chromosomal location of the 56 mutations detected in irradiated rice. Rice chromosomes are shown as boxes in blue with chromosomal numbers at the top and centromeres represented by yellow boxes. The mutation sites are indicated by horizontal bars in the chromosome boxes with the mutation number (left) and type (right) color-coded by the mutant lines: “3098” (red), “786-5” (blue), “885” (orange), “IRB 3517-3” (magenta) and “IRB 3790-2” (green). Numbers in parentheses indicate the size (bp) of the deletion or insertion. The green box in the upper arm of chromosome 6 shows the 535-kb inverted region of mutation #52. The position of centromeres is based on the Os-Nipponbare-Reference-IRGSP-1.0 database (RAP-DB https://rapdb.dna.affrc.go.jp/).

We compared the estimated MFs to previous genome-wide mutation studies in rice. The whole-exome sequencing of unselected M2 plants obtained from dry rice seeds irradiated with 150 Gy carbon ions (135 MeV/u, LET: 30 keV/μm) in RNC, RIKEN reported values for the number of mutations that were comparable to the present study, which was 9.06 ± 0.37 (average ± standard error) per plant [27]. The MF in M2 was 2.2 ×10^-7^ per base, which was estimated by dividing the number of mutations by the size of the enriched target region (41.8 Mb). The previously reported work utilized different ^12^C^6+^ ion beams (LET: 30 keV/μm), in terms of their physical properties, from that used in our study (LET: 76 keV/μm), therefore the selected doses are different (150 Gy in the previous work and 40 Gy in our study). In another case, whole genome sequencing analysis of M6 lines from “Hitomebore” rice seeds irradiated with 30 Gy ^12^C^5+^ ions (220 MeV, LET: 107 keV/μm) estimated an MF in M2 of 2.4 ×10^-7^ per base [28]. The exome sequencing analysis of EMS-mutagenized DNA with 39-Mb target probes reported that an MF in M2 plants was 5.2 × 10^-6^ per base [32]. Whole genome sequencing analysis of three independent tissue culture-regenerated rice plants suggested that 2,492, 1,039, and 450 SNVs occurred in the M1 generation, indicating that the average of MF for base substitution in M1 was 1.7 × 10^-6^ per base [33]. Another whole genome sequencing analysis of regenerated rice plants that were selfed for eight consecutive generations detected 54,268 DNA polymorphisms, including 37,332 SNPs and 16,936 small INSs and DELs, which indicated that the MF was 1.5×10^−4^ per site for all polymorphisms [34]. Conversely, large-scale whole genome sequencing analysis of a mixed M2 and M3 population of fast neutron (FN) “Kitaake” rice mutants identified 91,513 mutations in 1,504 lines, estimating the MF against the reference genome size (374 Mb) in this population to be 1.6 × 10^-7^ per base [35]. Whole genome sequencing analysis of M6 lines from “Hitomebore” rice seeds irradiated with 250 Gy gamma-rays estimated that the MF in M2 was 3.2 ×10^-7^ per base [30]. The number of detectable mutations with genome sequencing is highly dependent on the experimental conditions of mutagenesis, quality and quantity of the massive parallel sequencing, and the bioinformatics pipeline for sequence analysis. Although an accurate comparison is difficult for the these reasons, the data published to date including our results suggest that carbon ion irradiation induces smaller number of mutations in rice than EMS- and tissue culture-based mutagenesis and that this number is closer to that of FN and gamma-ray mutagenesis.

MFs of 3.4 × 10^−7^ single base mutation per base were also reported in carbon ion-mutagenized Arabidopsis M1 seeds exposed to 200Gy ^12^C^6+^ ions (43 MeV/u; average LET within samples, 50 keV/μm) [19]. Another report of Arabidopsis using 17.3 MeV/u carbon ions (surface LET, 107 keV/μm) at TIARA also showed MFs of 2.7 × 10^−7^ and 3.2 × 10^−7^ per base in M2 plants generated by dry seed irradiation (125 and 175 Gy, respectively), and 1.8 × 10^−7^ and 1.6 × 10^−7^ per base in M2 plants generated by seedling irradiation (20 and 30 Gy, respectively) [21]. Kazama et al. [20] reported 307 and 473 mutations that included rearrangements, single base substitutions, and small INSs and DELs in eight lines of argon ion- (290 keV/μm, 50 Gy) and carbon (30.0 keV/μm, 400 Gy) ion-mutagenized M3 populations obtained by dry seed irradiation in Arabidopsis, indicating MFs calculated with 119.7 Mb genome are 3.2 × 10^−7^ and 5.0 × 10^−7^ per base, respectively. Therefore, MFs caused by ion-beam irradiation seem to be the same in order of magnitude between Arabidopsis and rice.

Among the 56 mutations detected, 24 (43%), 23 (41%) and 5 (9%) mutations were SNVs, DELs, and INSs, respectively. The ratio of SNVs in this work was slightly lower than the irradiation results of a lower LET level of ^12^C^6+^ ions (LET 30 keV/μm) [27] in which more than half (58%) of the detected mutation were SNVs, followed by DELs (37%) and INSs (4%) in 110 independent M2 lines from dry rice seeds irradiated with 150 Gy of carbon ions. The FN mutagenesis of rice seeds also induced SNVs most frequently (48%) with DELs accounting for 35% [35]. While, mutagenesis with 30 Gy ^12^C^5+^ ions (220 MeV, LET: 107 keV/μm) or gamma-rays induced high ratio of SNVs, approximately 70 % of the detected mutations [28].

The most frequent base changes (38% of the total SNVs) were G to A and C to T (GC **→** AT) transitions (Fig 5A). Additionally, AT **→** GC transition accounted for 33% of the total SNVs and various types of base changes were induced. The proportion of base change pattern was similar to that in other ion beam-, gamma-ray- and FN-mutagenized rice [27,28,36] and in contrast to EMS-mutagenized rice, in which most of the mutations were SNVs of GC **→** AT transitions [32]. The ability to induce various types of base changes other than GC → AT transitions could be an advantage for effective mutagenesis in DNA regions or genomes with low GC content. In addition, ion-beam mutagenesis can cause 170 possible types of amino acid conversion achieved by single nucleotide mutations, while EMS requires G or C at nonsynonymous positions in amino acid codons and causes only 26 amino acid conversion types [37].

**Fig. 5.**
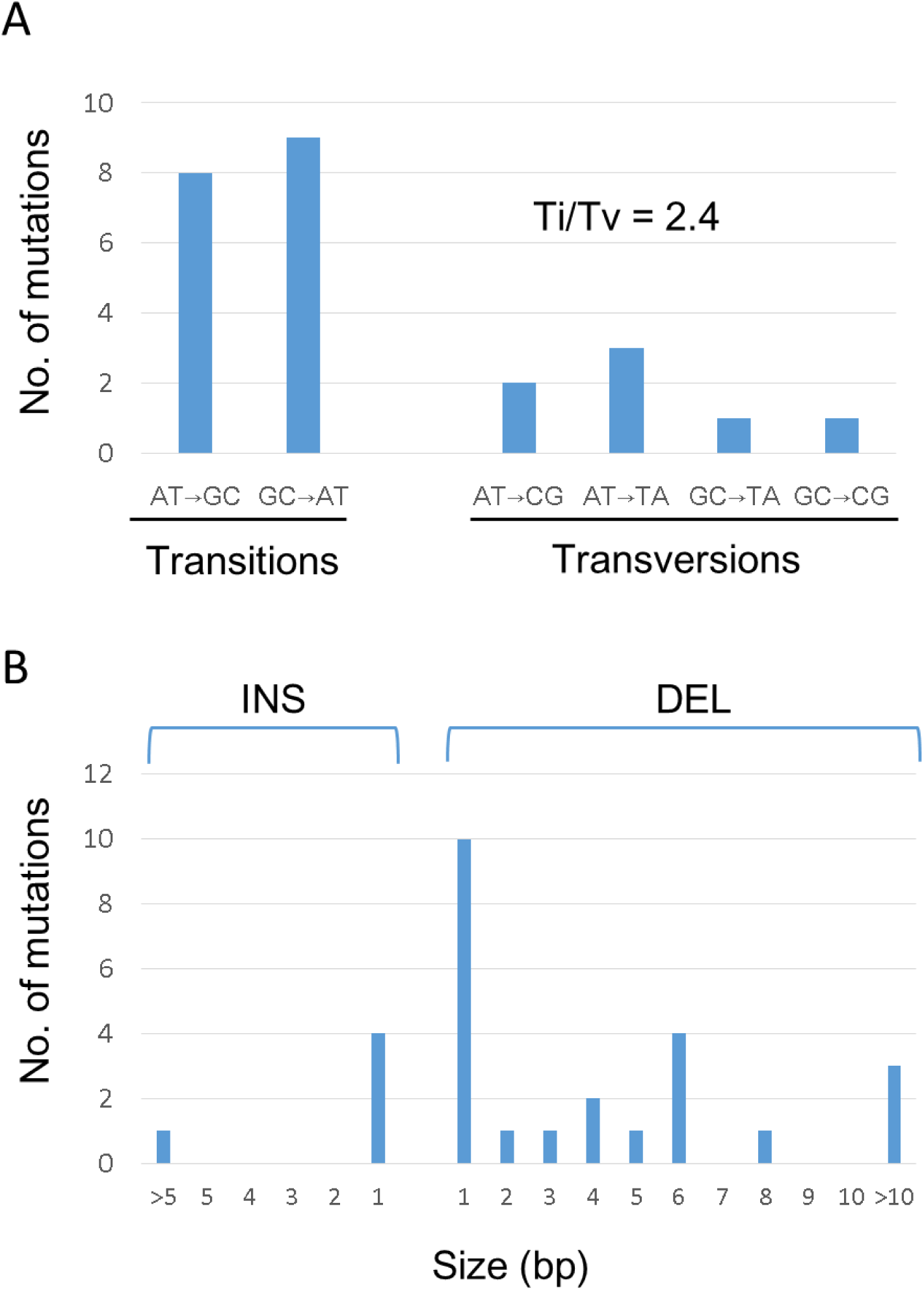
Summary of small mutations detected in rice mutagenized with ion-beam irradiation. (A) Base changes in 24 SNVs. Complementary mutations, such as A→C and T→G, are pooled. The ratio of transition to transversion (Ti/Tv) was 2.4. (B) Size distributions of 5 insertions (INS) and 23 deletions (DEL).

Of the 28 INSs and DELs (InDels), 14 (50%) were 1 bp in size and most of the InDels (89%) were less than 10 bp in length, excluding three relatively large DELs (mutations #7, #23, and #38) that were 128, 46, and 33.6 kb in length, respectively (Table 4 and Fig. 5B). Sixteen (57%) occurred within nucleotide repeats such as homopolymers that consisted of three or more repeats of the same nucleotides, or polynucleotide repeats that consisted of either partial or full repetitive sequences (Table 4). Many of the single-nucleotide InDels (eight of 14 events; 57%) were associated with homopolymers, but none of them were associated with a polynucleotide repeat. In contrast, InDels that were more than 2 bp in length tended to be localized near polynucleotide repeats (seven of 14 events; 50%) than homopolymers (only one of 14 events; 7.1%). No such repeat was observed in the DEL sites that were more than 7 bp in length. A similar pattern was observed in the presence of homopolymers or polynucleotide repeats at flanking sequences of small InDels in carbon ion-beam-irradiated Arabidopsis genome [19, 21]. Hase et al. [21] hypothesized that distinct mechanisms are involved between the generation of the single base deletion and larger-sized deletions in carbon ion-irradiated Arabidopsis. The authors also suggest that much larger sized deletions (equal or greater than 50 bp) are generated through another distinct mechanism. The presence or absence of characteristic repeats in the InDel sites in our data may also suggest that InDel generation mechanisms are different depending on the InDel size in the irradiated rice genome.

**Table 4.**
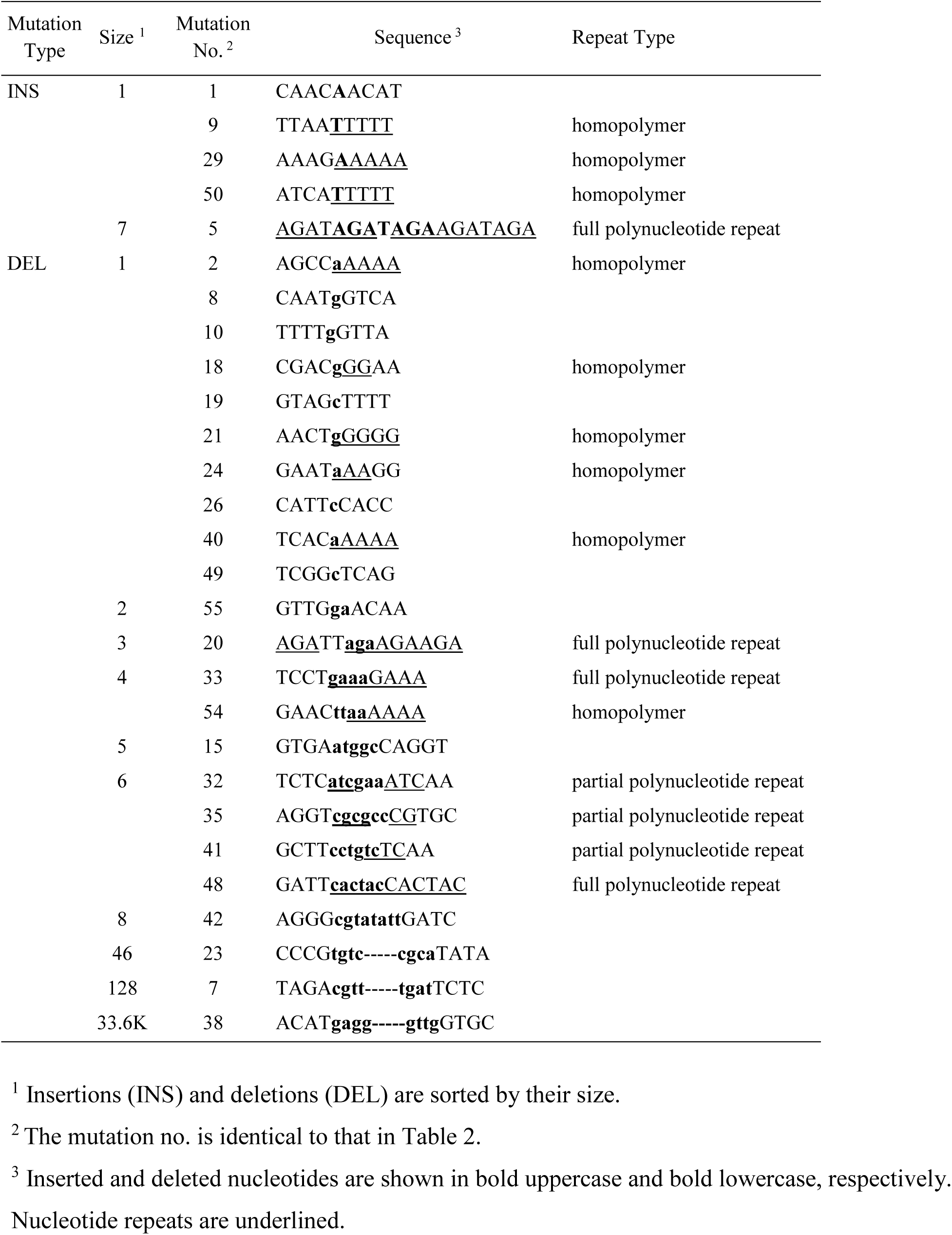
List of InDELs and repeat sequences

The primary structure of three mutations (#12, 13, and 37) classified as RPL are shown in Fig. 3. In two RPLs, #12 and #37, three and eight nucleotides were replaced with single bases accompanied by 2-bp and 7-bp deletions, respectively. Mutation #13 was a dinucleotide substitution in which GG was substituted with TA. A dinucleotide repeat, was found in the mutation site of #12 and #37 RPLs. In the 535-kb INV (mutation #52), 2-bp overlapping nucleotides were observed in one of the two junction sites, along with a 3-bp deletion on both sides of the mutated region (Fig 3). Although these mutation patterns were observed in RPLs, it was difficult to determine whether these features in primary sequences were related to DNA repair processes and mutation generation because the number of observed events was too small.

The effect of mutations on gene function was assessed with SnpEff [29] and we identified five and 13 mutations having putatively high (frameshift_variant and exon_loss_variant) and moderate (missense_variant and disruptive_inframe_deletion) impacts on protein functions classified according to the SnpEff output (S2 Table). In addition, by manually confirming the break points of an inversion (mutation #52), we found that one of the two breakpoints of the inversion was located in a coding region of the gene, *Os06g0275000* and thus, categorized as a high impact mutation. In total, six mutations were classified as high impact mutations with an average of 1.2 such high-impact mutations per line (Tables 2 and 3). Among these, an exon_loss_variant (mutation #38) consisted of a 33.6 kb deletion that caused a loss of five genes located within this genomic region. Thus, the number of affected genes located by six potentially high impact mutations was 10, with an average of two genes affected in each mutated line. Because high impact mutations have a higher probability of changing protein function, it is likely that they cause phonotypes observed in the mutants. Indeed, we found strong candidate mutations that potentially contribute to mutant phenotypes in four out of the five lines investigated.

### 3.3 Candidate genes that are responsible for mutant phenotypes

We isolated two dwarf mutants (lines “3098” and “885”) and three early-heading-date mutants (lines “786-5”, “IRB3517-3”, and “IRB3790-2”) in the present study (Fig. 2 and S1-S3 Figs). Using whole-exome sequencing analysis, we successfully identified candidate genes that are presumably responsible for the phenotypes observed in these mutants.

The mutant line “3098” was isolated as a dwarf mutant in the M2 population grown in the green house. Homozygosity of the dwarf phenotype was confirmed in M3 progenies grown in the green house as well as in the paddy field (Fig. 2A and S1 Fig. A, B and D). The panicles of line “3098” were compact and seeds were small and rounded compared with those of NPB (S1 Fig. E - H). The whole-exome sequencing analysis of DNA extracted from an M2 plant from line “3098” identified 10 mutations (Table 2 #1 –10 that included a 128-bp DEL (mutation #7) spanning the third intron and forth exon of the guanine nucleotide-binding protein alpha-1 subunit gene (*GPA1*, also called *RGA1*, and *D1*) that was predicted to cause a truncated protein (Fig. 6A). The functional disruption of this gene (*Os05g0333200*) causes the Daikoku dwarf (*d1*), which is characterized by its broad, dark green leaves, compact panicles, and small, round grains [38, 39]. These documented mutant phenotypes were apparent and quite similar to line “3098”. Although the initial analysis of whole-exome sequences suggested that this mutation (mutation #7) was present as a heterozygote in the M2 plant, PCR analysis using a specific primer set indicated that the mutation was present as a homozygote in the same M2 plant. PCR analysis of 12 F2 plants (six dwarf and six wild-type plants) from a backcross between line “3098” and NPB and subsequent self-crossing showed that all six dwarf plants harbored the 128-bp DEL (mutation #7) in homozygotes, whereas the remaining 6 were heterozygotes or wild type. Therefore, the 128-bp DEL in the *D1* gene was a strong candidate for the dwarf phenotype of line “3098”. The discrepancy in zygosity in the results from the initial bioinformatics call (heterozygous) and PCR (homozygous) as well as the phenotype (homozygous) could be due to the presence of nearly identical DNA sequence in the region of the *D1–like* gene (*Os05g0341300*), which is likely a pseudo gene on the same chromosome in the NPB genome [40]. A BLAST search of the 350-bp DNA sequence of the *D1* gene, including a 128-bp deletion and ∼100-bp upstream/downstream sequences (positions 15,612,701-15,613,050 on chromosome 5), indicated 99% identity with only a three-nucleotide difference in 350 bp within the region between 16,009,580 – 16,009,929 on chromosome 5, where the *D1–like* gene is located (Fig. 7). The nucleotide sequences were completely identical between the *D1* and *D1-like* genes in the 427-base upstream region of the deletion. Therefore, it was impossible to distinguish the origin of the short sequence reads from these regions, with some sequencing reads originating from the *D1–like* gene region were mapped to the deleted region in the *D1* gene. Consequently, the mutation was predicted to be heterozygous. Further improvement of the pipe line and mapping parameters for mutant calls may help to discriminate between very similar sequences.

**Fig 6.**
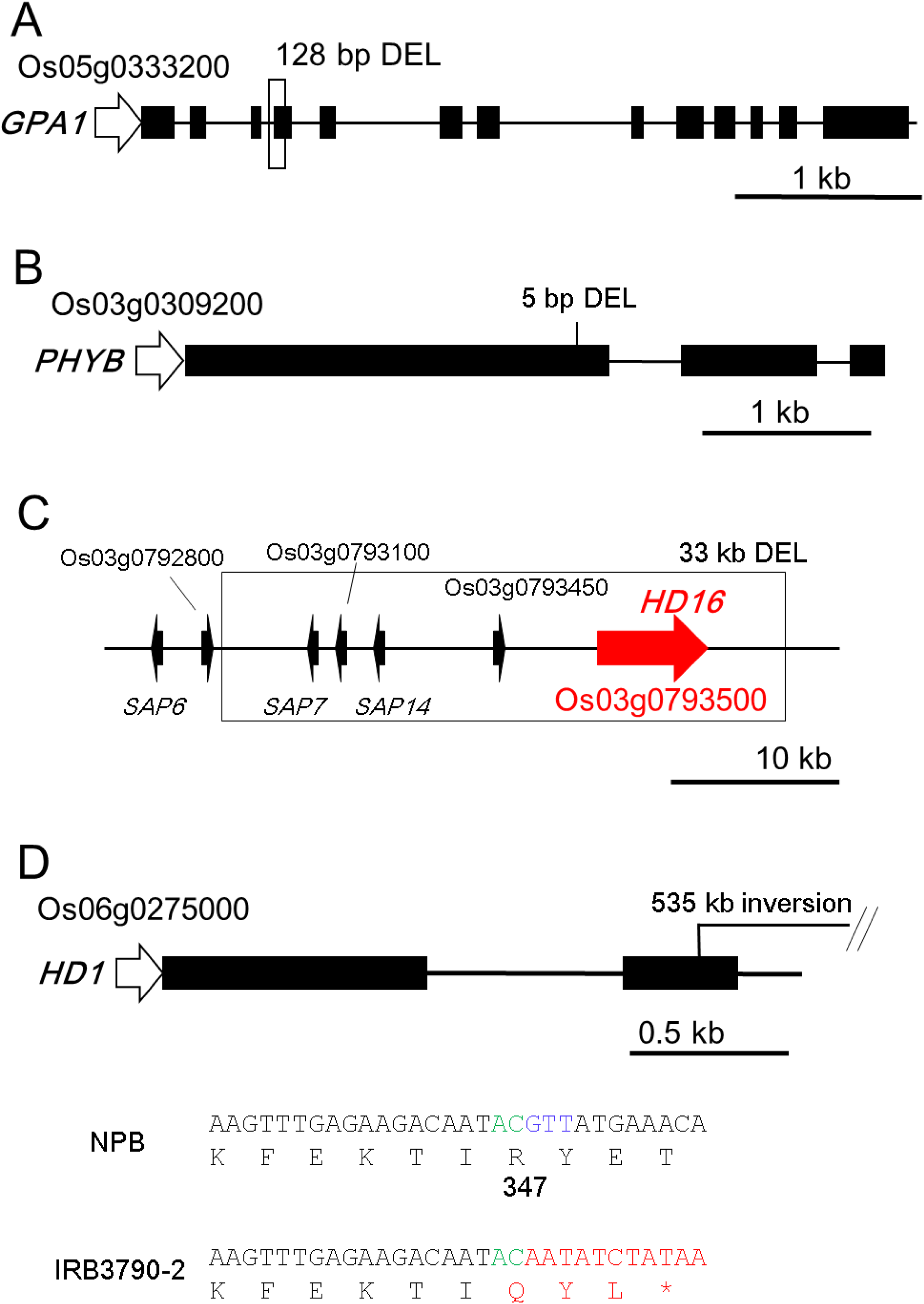
Structure of candidate genes and position of mutations in ion-beam irradiated rice. Black boxes represent exons, and a white arrow indicates the 5’ untranslated region. Introns and 3’ untranslated regions are indicated by black lines. (A) Guanine nucleotide-binding protein alpha-1 subunit (*GPA1*) gene. The position of mutation #7 in “3098”, the 128-bp DEL, indicated by an unfilled box. (B) Mutation #15 is the 5-bp DEL located in the first exon of the *PHYTOCHROM B* gene in the “786-5” line. (C) Relative position of the genes (solid arrows) in and near the 33-kb DEL (mutation #38) on chromosome 3 in the “IRB3517-3” line. The *HD16* gene is indicated in red. (D) One of the break points in the 535-kb INV of mutation #52 in “IRB3790-2” is located in the second exon of the *HD1* gene. Nucleotide and corresponding amino acid sequences of the break point in NPB control and “IRB3790-2” DNA are shown at the bottom of the panel. The amino acid (R347) of the HD1 protein is marked with the amino acid position number. The green and blue letters indicate overlapped and deleted nucleotides in junction sites. Inverted sequences, excluding overlapped nucleotides and altered amino acids, are shown in red. The detailed sequence of the junction sites is shown in Fig. 3.

**Fig 7.**
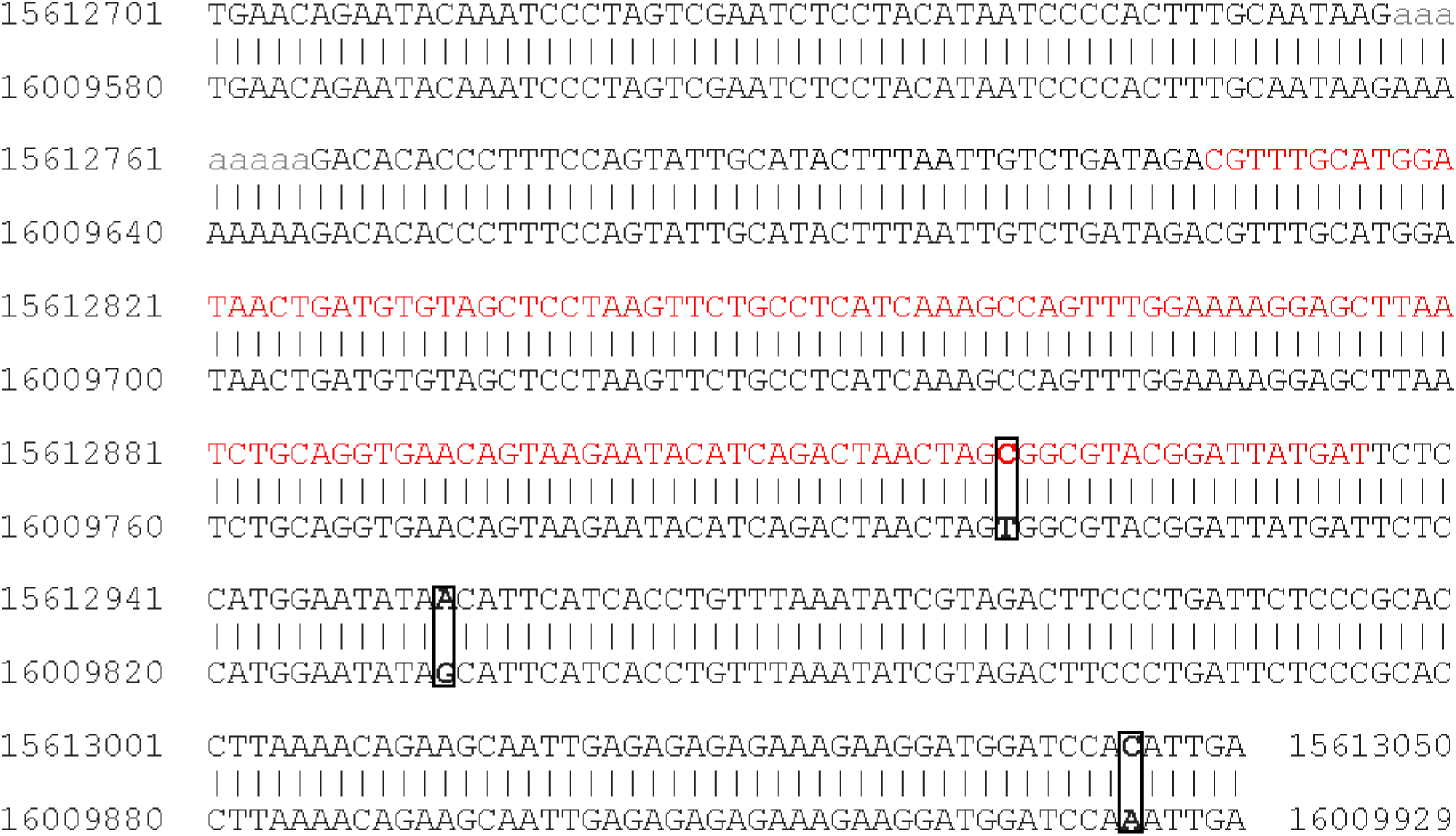
Alignment of the D1 and D1-like gene in rice. Alignment of the 350-bp *D1* gene region, including a 128-bp-deleted region (mutation #7 indicated) indicated in red letters plus ∼100-bp upstream/downstream sequences (Oschr05 15612701-15613050) and the similar *D1–like* gene region (Oschr05 16009580-16009929). Distinct nucleotides are indicated in bold letters surrounded by boxes. The numbers indicate the nucleotide position on chromosome 5.

Line “885”is a mutant line characterized by dwarfism and shoot bending (Fig. 2B). It was isolated from the M2 population grown in the greenhouse and its genetic homozygosity was confirmed in the M3 population in both the green house and paddy field (Fig. 2B and S1 Fig. C and D). Sixteen mutations were identified by whole-exome sequencing of genomic DNA from a single M3 plant (#20 - 35 in Table 2), but there was no potentially high impact mutation observed (Table3). Therefore, three homozygous nonsynonymous mutations (#30, #34, and #35) were selected to clarify whether these mutations were linked to the dwarf phenotype. PCR fragments amplified using primer sets for mutations #30, #34, and #35 (S1 Table) and template DNA extracted from four independent F2 plants with the dwarf mutant phenotype were subjected to Sanger sequencing. If the candidate genes are the causative genes, only mutated sequences would be detected in the PCR fragments from all four F2 plants. However, non-mutated sequences were detected in at least one of the F2 plants in all three mutation regions, suggesting that none of these three mutations were linked to the dwarf phenotype observed in the “885” line. It is possible that a mutation located outside of an exon region, presumably in the promoter, could be responsible for the dwarf phenotype observed in this line. Alternatively, the causative mutation could be located in a coding region but not effectively captured during library preparation or due to problem in the design of the capturing probe library. It may be also possible that the bioinformatics pipeline did not find the mutation due to complex sequence organization. Thus, an alternative approach such as whole genome sequencing and/or using a different bioinformatics pipe line is required to identify the causative mutation for this line.

The mutant line “786-5” was initially identified as a mutant candidate with wide leaves in the greenhouse (Fig 2C). Analysis of seedling morphology revealed that, in addition to wider second leaf blades, the length of second leaf blades was significantly shorter than that of the NPB control (S2 Fig. A - D). Furthermore, the mutant seedlings developed a third leaf earlier than NPB (S2 Fig. A and B) and had an increased angle between the second and third leaf blades (S2 Fig. E and F). The homozygosity for this line was confirmed in the M3 generation. In the paddy field; all M4 plants (n = 45) started heading 15 to 17 d earlier than the NPB control and showed broken flag leaves (Fig 2C and D). The whole-exome sequencing analysis of DNA extracted from an M3 plant identified nine mutations (#11 -19 in Table 2). Among these mutations, a 5-bp DEL (#15) was found in the first exon of the *PHYTOCHROME B* gene (*PHYB*; *Os03g0309200*), which caused a frameshift and was present in the homozygote (Fig 6B and Table 2). We considered this mutation to be the most likely candidate for the mutant morphology and early heading date phenotype in line “786-5” because similar phenotypes have also been reported in a rice *phyB* mutant [41].

The early-heading-date mutant “IRB3517-3” was identified in the M2 population. All M3 plants (n = 45) grown in the patty field showed earlier heading than the NPB control (Fig 2D), indicating that the genotype in the M2 plant was homozygous. The whole-exome sequencing analysis of DNA extracted from an M3 plants revealed 13 mutations (#36-48), including eight homozygous mutations (Table 2). The only high impact mutation identified was a 33.6-kb deletion on chromosome 3 (#38). The deleted region contained five genes, *Os03g0793000*, *Os03g0793100*, *Os03g0793300*, *Os03g0793450*, and *Os03g0793500* (Fig 6C), among which the *HEADING DATE 16* (*HD16*; *Os03g0793500*) gene was the most likely candidate for the early heading phenotype observed in the “IRB3517-3” mutant. The *HD16* gene encodes a casein kinase I protein and is known to act as an inhibitor of rice flowering by phosphorylating and activating the GHD7 protein, which one of the major players in the photoperiodic control of rice flowering [42]. It has been reported that a near-isogenic line with decreased kinase activity of the HD16 protein shows early flowering under a natural day length and late flowering under short-day conditions [42]. The late heading date phenotype under short-day condition was confirmed by growing the “IRB3517-3” mutant in a plant growth chamber (S3 Fig. A).

In another early-heading-date mutant line “IRB3790-2”, DNA from a single M3 plant obtained from a M2 plant that exhibited an early heading date phenotype was subjected to whole-exome sequencing analysis. However, this M2 plant was not homozygous for the early-heading-date phenotype, indicated by plants with early and normal heading date phenotypes that segregated in the M3 population (Fig 2D). We failed to establish a homozygous population even in the M4 generation and had to wait until M5 to obtain a homozygous population for this mutant line (S3 Fig. B). The sequencing results indicated that there were eight mutations (#49-56), including one high impact mutation (#52) that was a 543-kb INV on chromosome 6 (Table 2). One of the break points of this INV was located in the second exon of the *HEADING DATE 1* (*HD1*; *Os06g0275000*) gene (Fig 6D), which encodes a zinc-finger type transcription factor that regulates the expression of HEADING DATE 3, a mobile flowering signal [43, 44]. Although the position of the break point (R347) is near the C-terminal region of the encoded protein, the truncated HD1 protein generated by the INV lacks the CCT motif (amino acid position from 326-369), a conserved motif that is widely present in flowering-related proteins and essential for the protein function of HD1 [45]. These facts suggest that the truncated HD1 protein produced in “IRB3790-2” was a loss-of-function protein. PCR analysis of eight selected early and 12 non-early-heading-date individuals from the M4 population (scored on August 10, 2018) in the paddy field using primers detecting the #52 INV mutation showed that all eight early heading date plants contained this INV in both alleles, while that other plants did not. Moreover, M5 “IRB3790-2” plants were homozygous for this INV mutation. These results suggest that this inversion is likely the cause of the early heading date phenotype. Functional HD1 inhibits heading under natural field conditions but promote heading in short-day conditions [43]. Similarly, the M5 “IRB3790-2” population showed late heading date in short-day conditions (S3 Fig. A). We did not determine why homozygous *hd1* plants were not obtained until the M4 generation even though we chose a plant that exhibited the earliest heading date phenotype in the population to obtain seeds for the next generation in the paddy field. Heterozygous plants (*HD1* / *hd1*) are thought to show an intermediate phenotype between wild-type and *hd1* homozygous plants [46]; therefore, one possibility is that an additional mutated gene that affects heading date or viability could also present in the genome of the plants selected for propagation. Further genetic analysis may reveal additional factors that contribute to heading date determination in this mutant line.

In summary, we have identified likely candidate genes responsible for mutant phenotypes in four of the five mutant lines examined. Although further analyses such as genetic complementation, RNA inactivation, and/or genome editing, are required to confirm the relationship between the candidate genes and observed mutant phenotypes, this work shows that the combination of ion-beam mutagenesis and whole-exome sequencing can quickly narrow down probable candidate genes in rice. Because the average number of high impact mutations per line was very low, it was easy to identify candidate genes for the mutant phenotypes observed even if the responsible gene was an uncharacterized gene by examining the genetic linkage between the identified mutation and the phenotype.

Although a recent improvement in the genetic resources for rice enabled us to take a reverse genetic approach to understand the genetic basis of gene function and improve economically important traits in rice, such as the use of transposon- and T-DNA-tagged mutant libraries [47] and genome editing techniques [48], a forward genetics approach with induced mutagenesis is still important for identifying mutants with uncharacterized phenotypes or genes that cause these phenotypes. Induced mutagenesis can be applied not only in rice but also in other non-model plants that do not fulfil genetic information and resources. It would be particularly useful to introduce new phenotypic variations into plants with a specific genetic background or to improve traits of already established varieties used in agricultural production. Unlike tagged transposon- or T-DNA mutations, induced mutagenesis has not been thought to be advantageous for isolating a causal gene because it requires large efforts such as positional cloning. However, recent “mapping by sequencing” techniques that combine mutagenesis techniques and massive parallel sequencing have been used and promoted for the rapid detection of causal genes in mutants isolated by forward genetics [49]. For example, in the MutMap method [50], one of the mapping by sequencing techniques developed to identify rice genes, selected EMS-generated mutants were back-crossed to obtain the F2 generation. Then, DNA from F2 individuals that expressed mutant phenotype was pooled, subjected to NGS, and SNPs that were commonly present in mutant F2 DNA were detected by scoring the SNP index. This technique can provide an accurate evaluation of mutant phenotypes because it uses mutants and their parental lines in almost identical genetic backgrounds. It also does not need to generate new polymorphic DNA markers for mapping because massive parallel sequencing can detect many SNPs in EMS-generated F2 populations, which dramatically shortens the time required to clone a gene. As we discussed, ion beams induce a relatively small number of mutations compared with EMS and more effectively cause InDels that lead to complete inactivation of gene function. Therefore, it can be expected that a small number of genes will be listed as candidate genes responsible for mutant phenotype by ion-beam irradiation compared to EMS treatment. In this work, we successfully identified candidate genes from four mutants in a very efficient manner by evaluating mutation libraries generated by whole-exome sequencing. Furthermore, in Arabidopsis, the combination of ion-beam mutagenesis, whole genome sequencing, and rough mapping also successfully identified a causal gene of a variegated leaf mutant phenotype [51]. Therefore, characterization of ion beam-induced mutants by whole-exome or whole-genome sequencing will enable the promotion of effective isolation of causal genes for mutants.

Although we chose whole-exome sequencing in this work, given the recent cost reduction in next generation sequencing and the enrichment of genome data, whole-genome sequencing may be more applicable in rice and other model crops. Long-read NGS technologies, such as Oxford nanopore technology and PacBio are also becoming common NGS methods [52]. Although our bioinformatics pipeline successfully detected large mutations spanning one or more exons by the short-read sequencing as shown in this work and others [27], it may be possible that large InDels and INVs as well as chromosomal translocations are more effectively detected by the long-read sequencing. The long-read sequencing could be a good option for detecting causal gene candidates. High error rates and high cost, however, are the main limitations of current long-read NGS technologies [52]. For example, a small InDel that could also be a high impact mutation can be detected only by accurate sequencing. Therefore, combining short-read and long-lead technologies could be the good approach for detecting such candidates. Further efforts to optimize NGS technologies that identify useful genes from ion-beam-induced mutants will facilitate the improvement of agronomically important traits in rice as well as in other plants.

## 3.4 Conclusions

The properties of mutations in rice induced by ^12^C^6+^ ion beams accelerated using the TIARA AVF cyclotron at TARRI were characterized by whole-exome sequencing. In total, 56 mutations were detected in the 5 selected mutant lines and the average mutation frequency in the M1 generation of ion beam-irradiated rice was estimated to be 2.7 × 10^-7^ per base. Of the 56 mutations detected, six mutations were classified as high impact mutations with an average of 1.2 such high-impact mutations per line. The identification of a small number of high-impact mutation would facilitate identification of candidate genes responsible for the mutant phenotypes.

## Acknowledgements

The authors thank Shoya Hirata, Eenzen Mungunchudur, Toshihiko Sanzen, Hiroki Arai, Shogo Ozawa, Satoshi Kitamura, and members of the Research Planning Office, Quantum Beam Research Directorate, QST for their help in harvesting rice seeds and characterizing the rice mutants, Dr. Feng Li from RBD, NICS, NARO for his valuable comments on the manuscript. The bioinformatics analysis was performed using the HOKUSAI supercomputing system, operated by the Information Systems Division, RIKEN, under the project number Q18208. This work was supported by a grant from the Cabinet Office, under the “Technologies for creating next-generation agriculture, forestry and fisheries” project in the Cross-ministerial Strategic Innovation Promotion Program (SIP). The grant was administrated by the Bio-oriented Technology Research Advancement Institution, NARO.

## Author contributions

Y.O and Y.H designed the experiment. S.N. and Y.H. conducted ion-beam irradiation, mutant screening, and characterization of mutants in the green house. Y.O., S.N., K.S., A.S., H.K., and Y.H. performed mutant screening and characterization of mutants in the paddy field. Whole exome analysis was done by H.I., R.M., and T.A.. Y.O. and H.I. analyzed the data and Y.O. wrote the manuscript with input from H.I. and Y.H. All authors approved the final manuscript.

## Supporting Information

**S1 Fig.**
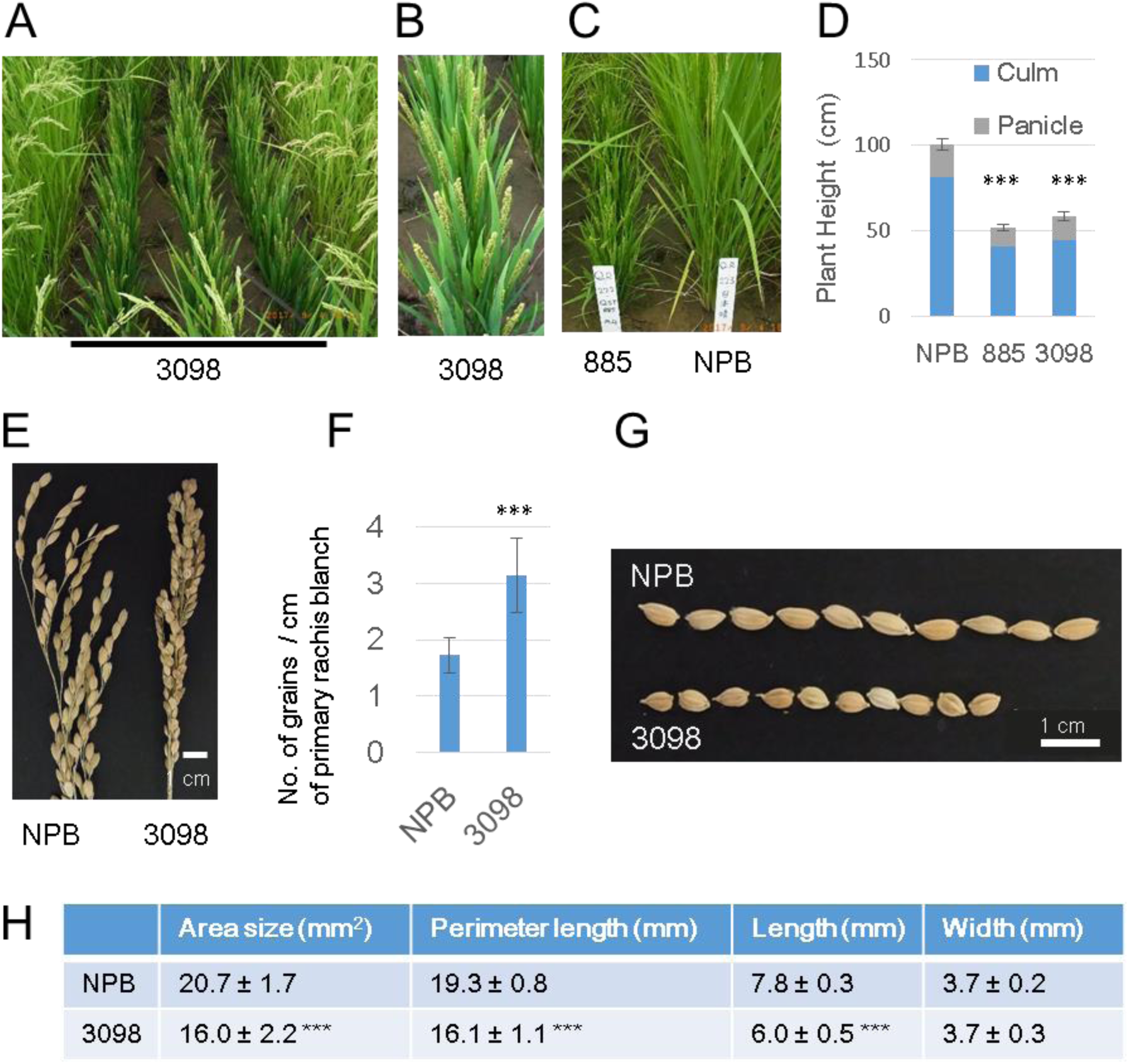
Phenotype of dwarf mutants grown in the paddy field. (A) The dwarf mutant line, “3098” (three middle lanes) and NPB plants (most left and right) grown in the paddy field. (B) A magnified photo of “3098” in A. (C) The dwarf mutant line, “885” (left) and NPB plants (right) grown in the paddy field. (D) Comparison of plant height (length of culms and panicles) among the two dwarf mutants (“885” (M4) and “3098” (M3)) and NPB control recorded in the paddy field. Error bars indicate SD (n = 8∼9). P values in two-sample *t*-tests were < 0.001 for both NPB vs “885” and NPB vs “3098”. For (A-D), one-month-old seedlings were transferred to the paddy field on May 24, 2017. The photograph was taken on September 4, 2017 and plant height was measured on October 5, 2017. (E)) A photograph of the panicles of NPB control and “3098”plants. (E) Comparison of grain number per length (cm) of a randomly selected panicle primary rachis blanch (n = 14). P value in two-sample *t*-tests was < 0.001. (F)) A photograph of the grains from NPB control and “3098”plants. (G) Comparison of the shape of the grains collected from NPB control and “3098”rice plants. Area, perimeter length, grain length, and width (mean ± SD, n = 50) were measured withy SmartGRAIN image analysis software obtained from http://phenotyping.image.coocan.jp/smartgrain/index.html [53]. Differences in area, perimeter length, and seed length were significant (***, P < 0.001) between NPB control and “3098”, whereas differences in grain width was not (P = 0.28).

**S2 Fig.**
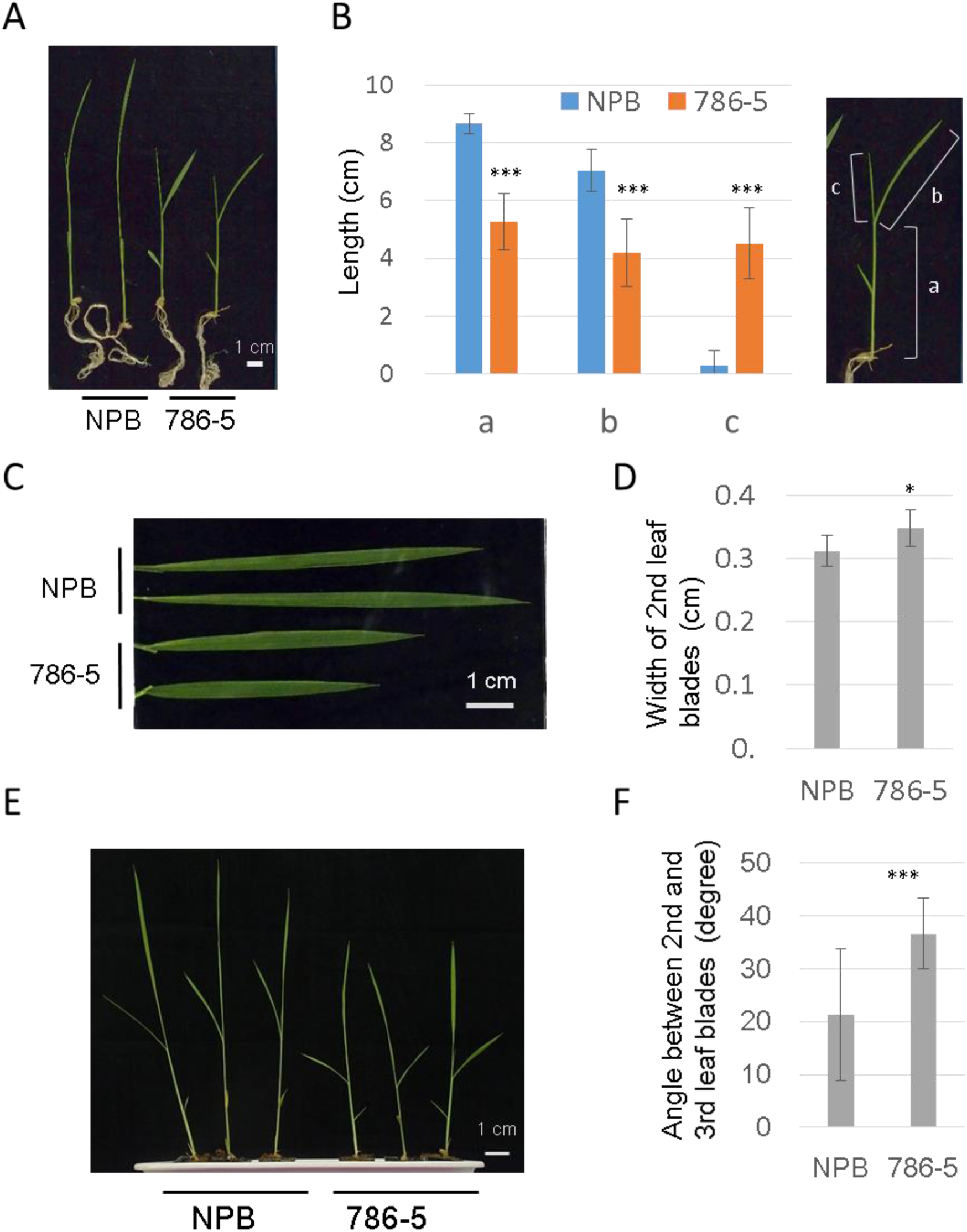
Phenotype of “786-5” rice seedlings. (A and B) A representative image of 14-d-old NPB and “786-5” (M5) rice seedlings grown in the green house (A) and the average distance from culm base (height) to lamina joint of second leaf (a) and length of second (b) and third (c) leaf blades (B). (C and D) A photograph of the second leaf blades on 14-d-old NPB and “786-5” seedlings (C) and the average width of the second leaf blades (E) that were measured with ImageJ software (n= 8). (E and F) A photograph of 20-d-old NPB and “786-5” rice seedlings grown in the green house (E) and the angles between second and third leaf blades (F) that were measured with ImageJ software (n= 10). Asterisks (*** and *) indicate that P values in a two-sample *t*-tests were < 0.001 and < 0.05 for NPB vs “786-5”, respectively.

**S3 Fig.**
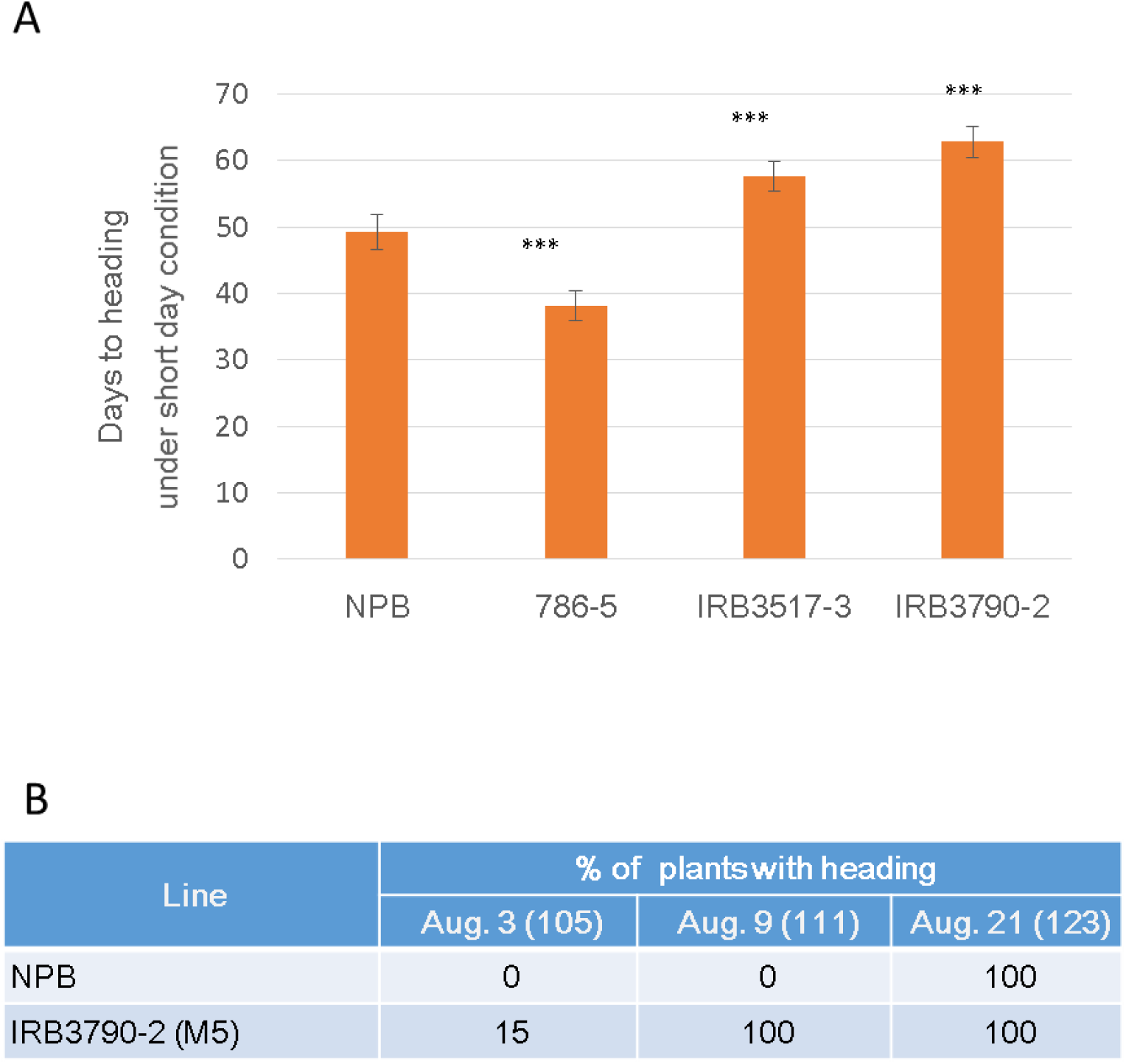
Additional data for heading date. (A) The heading dates of NPB and “IRB3790-2” were recorded in the paddy field in 2018. The M5 population was used for “IRB3790-2”. Approximately one-month-old seedlings were transferred to the paddy field on May 24, 2018. Ratios (%) of plants with heading recorded on August 3, 9, and 21 are shown. Numbers in parentheses indicate days after sowing. (B) Days to heading under short day light condition (10-h light / 14-h dark). Rice plants were grown under florescent lamps at 28°C in a plant growth chamber (BIOTRON LH200, NKsystem, Tokyo Japan). M5, M4, and M6 plants were tested for “786-5”, “IRB3517-3”, and “IRB-3790-2”, respectively. Asterisks (***) indicate P values < 0.001 in a two-sample *t*-test between NPB (49.2 ± 2.6 days, n =10) and the mutant lines, “786-5” (38.1 ± 2.2 days, n = 8), “IRB3517-3” (57.6 ± 2.3 days, n = 10), and “IRB-3790-2” (62.8 ± 2.3 days, n = 10).

S1 Table. List of primer sets

S2 Table. Detailed list of mutations detected

## Notes

#### Summary of Updates

All parts of the manuscript are updated to clarify. Figure 2 is revised and supplemental figures are added.

