## Supplementary material for "Genome sequencing of ion-beam-induced mutants facilitates detection of candidate genes responsible for phenotypes of mutants in rice": S1 Table

**S1 Table. List of primer sets**

| Mutation No.[1] | Line | CHR No. | Target Start | Target End | Target Reference[2] | Target Alternatives[2] | Type of mutation[3] | 5' position of primer[4] | Sequence (5' to 3') of primer |
|---|---|---|---|---|---|---|---|---|---|
| 5 | 3098 | chr04 | 2.3E+07 | 22824954 | T | TAGATAGA | INS (7) | chr4 22824682 F | GGCAAACATTACATATAGCATAAAGG |
| | | | | | | | | chr4 22825181 R | GAACTCGCCGACCTCACC |
| 7 | 3098 | chr05 | 1.6E+07 | 15612936 | ACGT….TGA | A | DEL (128) | chr5 15612143 F | GAACAAGGTAAATGCACAAAGGTATA |
| | | | | | | | | chr5 15613069 R | TTTTAGTTTGAACTTGTTCTCAATG |
| 12 | 786 | chr01 | 4.2E+07 | 41549064 | GTGT | GA | RPL | chr1 41548923 F | CTGCGCTCTTCAACCGTTC |
| | | | | | | | | chr1 41549206 R | CGGCGTGTTATTGCTCAGT |
| 13 | 786 | chr03 | 1631231 | 1631232 | GG | TA | RPL | chr3 1630969 F | CACTGCTCGGAGAACACCTT |
| | | | | | | | | chr3 1631462 R | TGGCTGGCTACGTGTGTAGA |
| 15 | 786 | chr03 | 1.1E+07 | 11022260 | AATGGC | A | DEL (5) | chr3 11022016 F | TCCGGTCACACACAGCTAAG |
| | | | | | | | | chr3 11022491 R | TGTTTCCTCAGATTCCTTGAAGA |
| 18 | 786 | chr10 | 2.1E+07 | 21181034 | CG | C | DEL (1) | chr10 21180767 F | GCTGTCCATCAGGTTCCTCT |
| | | | | | | | | chr10 21181262 R | ATTCGATGGATGGCTAGCTG |
| 22 | 885 | chr01 | 3357475 | 3357475 | C | T | SNV | chr1 3357244 F | GCGCAACATGAAAAAGAAAA |
| | | | | | | | | chr1 3357725 R | CCAGCGCAGTAGGAAATGAT |
| 23 | 885 | chr01 | 3.8E+07 | 37524065 | GTGT….CGC | G | DEL (46) | chr1 37523767 F | GCGCATGTTTGTGAGAGAAG |
| | | | | | | | | chr1 37524250 R | CGAAGCGAGCAATCTCAAGT |
| 26 | 885 | chr03 | 3.6E+07 | 35501011 | TC | T | DEL (1) | chr3 35500787 F | TGTTATCAGGGGCCTATTGAA |
| | | | | | | | | chr3 35501282 R | CATGACAGATGATTGGAACTTTG |
| 27 | 885 | chr04 | 2.4E+07 | 23792560 | G | A | SNV | chr4 23792336 F | TTTCTGATCTCGCCTTGGTT |
| | | | | | | | | chr4 23792789 R | ACAGGAGCAAGAACCGGAAG |
| 30 | 885 | chr05 | 2.2E+07 | 21895103 | C | T | SNV | chr5 21894866 F | CCAGTGGGATTTCCTCTGAA |
| | | | | | | | | chr5 21895323 R | CAAAAAGCAAATGTCCACGA |
| 32 | 885 | chr08 | 1.4E+07 | 14099489 | CATCGAA | C | DEL (6) | chr8 14099248 F | GTAAGCGCATGGAAATACCG |
| | | | | | | | | chr8 14099767 R | GGACACGGCTGAGAGGAG |
| 33 | 885 | chr08 | 2.5E+07 | 24691425 | TGAAA | T | DEL (4) | chr8 24691196 F | GCTAAATTTTAAGCACCATGTGA |
| | | | | | | | | chr8 24691653 R | CTGCTCCAACTCTGCATCAA |
| 34 | 885 | chr08 | 2.8E+07 | 27686683 | T | C | SNV | chr8 27686445 F | TTGCCTTCTAGGCCTCACTC |
| | | | | | | | | chr8 27686938 R | TGCTTTGACAGGAGATGCAG |
| 35 | 885 | chr10 | 1.6E+07 | 15810767 | TCGCGCC | T | DEL (6) | chr10 15810518 F | CACCGGAACGACCTCACC |
| | | | | | | | | chr10 15810983 R | CTCGCAATACCATGGAGGAC |
| 37 | IRB3517-3 | chr01 | 3.3E+07 | 32645636 | ACTCTGCAT | AA | RPL | chr1 32645404 F | TCTGACAAAAATAGCCCCAAA |
| | | | | | | | | chr1 32645884 R | CCATGTTGAAAGGAAACCAGA |
| 38 | IRB3517-3 | chr03 | 3.3E+07 | 33010771 | TGAG….GTT | T | DEL (33.6 k) | chr3 32976920 F | TGGCGTCGGGGTTTGTAT |
| | | | | | | | | chr3 33011012 R | AACACCACCACAAAGGAAGC |
| 41 | IRB3517-3 | chr06 | 2.8E+07 | 27521464 | TCCTGTC | T | DEL (6) | chr6 27521228 F | CCAGGTCACAACCATGTCAA |
| | | | | | | | | chr6 27521682 R | GCTGGTTTTGAAGCTGCATT |
| 42 | IRB3517-3 | chr07 | 4881911 | 4881919 | GCGTATATT | G | DEL (8) | chr7 4881645 F | CACGACGCCGAGAAGTCTAT |
| | | | | | | | | chr7 4882141 R | AAGCCCCTACAAAAGCCAAG |
| 44 | IRB3517-3 | chr09 | 7224224 | 7224224 | A | G | SNV | chr9 7223954 F | CCGAGGAAAAGTGCTGAAAG |
| | | | | | | | | chr9 7224446 R | GGGATGGAAGTTTGTTTTCG |
| 45 | IRB3517-3 | chr09 | 1.3E+07 | 13007477 | G | A | SNV | chr9 13007214 F | CCTGACCACCAAAAGAAAGG |
| | | | | | | | | chr9 13007708 R | ACGAGGACCTGCTGACAGA |
| 47 | IRB3517-3 | chr11 | 7365282 | 7365282 | T | C | SNV | chr11 7365028 F | ACCCATCAGCTGGTCACAAT |
| | | | | | | | | chr11 7365527 R | TTTGCTGTTTGTATGCTTCATTT |
| 52 | IRB3790-2 | chr06 | 9338213 | 9872884 | TATG….TTG | TACA….TCA | INV (535 k)[5] | chr6 9338078 F | CCTGCTGGAGCAATCAATCT |
| | | | | | | | | chr6 9338377 R | AGCGTCTCATGAGTCCCATC |
| | | | | | | | | chr6 9872652 F | CACTCATTAACAGAAGGGACTGAA |
| | | | | | | | | chr6 9872951 R | GTACTCCCTCCGTCCGAAA |
| 54 | IRB3790-2 | chr08 | 2.2E+07 | 21594853 | CTTAA | C | DEL (4) | chr8 21594571 F | CCAGTTTCATCCCAAAAATGA |
| | | | | | | | | chr8 21595070 R | TTTGTGTGTGACATATGAACGA |
| 56 | IRB3790-2 | chr12 | 4637905 | 4637905 | G | A | SNV | chr12 4637679 F | ACAAAAGCTGCAGCAGGAGA |
| | | | | | | | | chr12 4638153 R | GTCGCTCCAAGGGTGGAA |

[1] Mutation No. is identical to the one in Table 2.

[2] For the large deletions and the inversion, only first and last 4 bases are shown.

[3] Size of InDel is indicated in parentheses.

[4] "F" and "R" indicate orientation of the primers, forward and reverse, respectively.

[5] DNA fragments can be amplified with the PCR primer sets chr6 9338078 F-chr6 9872652 F and chr6 9338377 R-chr6 9872951R for the INV mutation.
