## Supplementary material for "Genome sequencing of ion-beam-induced mutants facilitates detection of candidate genes responsible for phenotypes of mutants in rice": S2 Table

**S2 Table. Detailed list of mutations detected**

| Mutation No.[1] | Line | CHR No. | Start | End | Reference [2] | Alternatives [2] | Type after evaluatio | Effect [4] | Gene | Annotation | Genotype[5] | Verification [6] |
|---|---|---|---|---|---|---|---|---|---|---|---|---|
| 1 | 3098 | chr02 | 16190324 | 16190325 | C | CA | INS (1) | intergenic_region | | | 0\|1 | |
| 2 | 3098 | chr02 | 25123707 | 25123708 | CA | C | DEL (1) | downstream_gene_variant | | | 0\|1 | |
| 3 | 3098 | chr03 | 17837150 | 17837150 | C | G | SNV | splice_region_variant&intron_variant | Os03g0426900 | [CLPB-C] Similar to APG6/CLPB-P/CLPB3 (ALBINO AND PALE GREEN 6); ATP binding / ATPase. | 0\|1 | |
| 4 | 3098 | chr03 | 21504174 | 21504174 | T | C | SNV | missense_variant | Os03g0583900 | [DCL2A] Similar to Endoribonuclease Dicer homolog 2a. | 0\|1 | |
| 5 | 3098 | chr04 | 22824947 | 22824954 | T | TAGATAGA | INS (7) | upstream_gene_variant | | | 1\|1 | ✓ |
| 6 | 3098 | chr05 | 5977789 | 5977789 | C | T | SNV | missense_variant | Os05g0196800 | Similar to Diacylglycerol acylCoA acyltransferase. | 0\|1 | |
| 7 | 3098 | chr05 | 15612808 | 15612936 | ACGT....TGAT | A | DEL (128) | frameshift_variant&splice_acceptor_variant&splice_region_variant&splice_region_variant&intron_variant | Os05g0333200 | [D1] Guanine nucleotide-binding protein alpha-1 subunit (GP-alpha-1). | 0\|1 | ✓ |
| 8 | 3098 | chr08 | 3299965 | 3299966 | TG | T | DEL (1) | downstream_gene_variant | | | 0\|1 | |
| 9 | 3098 | chr08 | 12412765 | 12412766 | A | AT | INS (1) | downstream_gene_variant | | | 0\|1 | |
| 10 | 3098 | chr12 | 17352209 | 17352210 | TG | T | DEL (1) | intergenic_region | | | 0\|1 | |
| 11 | 786 | chr01 | 25949685 | 25949685 | T | A | SNV | missense_variant | Os01g0644600 | Glutelin family protein. | 0\|1 | |
| 12 [7] | 786 | chr01 | 41549062 | 41549064 | GTGT | GA | RPL | upstream_gene_variant | | | 1\|1 | ✓ |
| 13 [7] | 786 | chr03 | 1631231 | 1631232 | GG | TA | RPL | missense_variant | Os03g0128800 | Similar to predicted protein. | 1\|1 | ✓ |
| 14 | 786 | chr03 | 2403133 | 2403133 | A | T | SNV | upstream_gene_variant | | | 1\|1 | |
| 15 | 786 | chr03 | 11022255 | 11022260 | AATGGC | A | DEL (5) | frameshift_variant | Os03g0309200 | [PHYB] Similar to Phytochrome B. | 1\|1 | ✓ |
| 16 | 786 | chr06 | 5399955 | 5399955 | A | C | SNV | intron_variant | | | 1\|1 | |
| 17 | 786 | chr07 | 13311076 | 13311076 | A | T | SNV | upstream_gene_variant | | | 1\|1 | |
| 18 | 786 | chr10 | 21181033 | 21181034 | CG | C | DEL (1) | frameshift_variant | Os10g0542400 | Expansin/Lol pI family protein. | 1\|1 | ✓ |
| 19 | 786 | chr12 | 23844768 | 23844769 | GC | G | DEL (1) | frameshift_variant | Os12g0577100 | Nucleotide-binding, alpha-beta plait domain containing protein. | 0\|1 | |
| 20 | 885 | chr01 | 1319914 | 1319917 | TAGA | T | DEL (3) | intergenic_region | | | 1\|1 | |
| 21 | 885 | chr01 | 3020438 | 3020439 | TG | T | DEL (1) | intergenic_region | | | 0\|1 | |
| 22 | 885 | chr01 | 3357475 | 3357475 | C | T | SNV | synonymous_variant | | | 1\|1 | ✓ |
| 23 | 885 | chr01 | 37524019 | 37524065 | GTGT….CGCA | G | DEL (46) | 5_prime_UTR_variant | Os01g0866500 | Similar to Soluble inorganic pyrophosphatase (EC 3.6.1.1) (Pyrophosphate phospho- hydrolase) | 1\|1 | ✓ |
| 24 | 885 | chr02 | 10379090 | 10379091 | TA | T | DEL (1) | 5_prime_UTR_variant | Os02g0280400 | Similar to casein kinase I isoform delta-like. ; Similar to Dual specificity kinase 1. | 0\|1 | |
| 25 | 885 | chr03 | 7473040 | 7473040 | T | C | SNV | upstream_gene_variant | | | 0\|1 | |
| 26 | 885 | chr03 | 35501010 | 35501011 | TC | T | DEL (1) | upstream_gene_variant | | | 1\|1 | ✓ |
| 27 | 885 | chr04 | 23792560 | 23792560 | G | A | SNV | upstream_gene_variant | | | 1\|1 | ✓ |
| 28 | 885 | chr05 | 318166 | 318166 | G | A | SNV | missense_variant | Os05g0105900 | Nucleotide-binding, alpha-beta plait domain containing protein. | 0\|1 | |
| 29 | 885 | chr05 | 15013747 | 15013748 | G | GA | INS (1) | intergenic_region | | | 0\|1 | |
| 30 | 885 | chr05 | 21895103 | 21895103 | C | T | SNV | missense_variant | Os05g0446600 | Similar to DNA glycosylase/lyase 701. | 1\|1 | ✓ |
| 31 | 885 | chr08 | 5808016 | 5808016 | T | C | SNV | downstream_gene_variant | | | 1\|1 | |
| 32 | 885 | chr08 | 14099483 | 14099489 | CATCGAA | C | DEL (6) | intergenic_region | | | 1\|1 | ✓ |
| 33 | 885 | chr08 | 24691421 | 24691425 | TGAAA | T | DEL (4) | downstream_gene_variant | | | 1\|1 | ✓ |
| 34 | 885 | chr08 | 27686683 | 27686683 | T | C | SNV | missense_variant | Os08g0553400 | Hypothetical conserved gene. | 1\|1 | ✓ |
| 35 | 885 | chr10 | 15810761 | 15810767 | TCGCGCC | T | DEL (6) | disruptive_inframe_deletion | Os10g0439924 | [CYP71Z8] Cytochrome P450 family protein. | 1\|1 | ✓ |
| 36 | IRB3517-3 | chr01 | 22075801 | 22075801 | G | T | SNV | missense_variant | Os01g0574400 | Similar to Cell division protein ftsH (EC 3.4.24.-). | 0\|1 | |
| 37 [7] | IRB3517-3 | chr01 | 32645629 | 32645636 | ACTCTGCAT | AA | RPL | intron_variant | | | 1\|1 | ✓ |
| 38 | IRB3517-3 | chr03 | 32977149 | 33010771 | TGAG….GTTG | T | DEL (33.6k) | exon_loss_variant&splice_region_variant | Os03g0793000 ~ Os03g0793500 [8] | 5 ORFs are deleted [8] | 1\|1 | ✓ |
| 39 | IRB3517-3 | chr04 | 26873179 | 26873179 | A | C | SNV | intergenic_region | | | 1\|1 | |
| 40 | IRB3517-3 | chr06 | 9657976 | 9657977 | CA | C | DEL (1) | intergenic_region | | | 0\|1 | |
| 41 | IRB3517-3 | chr06 | 27521458 | 27521464 | TCCTGTC | T | DEL (6) | disruptive_inframe_deletion | Os06g0665800 | [HMA9] Similar to heavy metal ATPase. | 1\|1 | ✓ |
| 42 | IRB3517-3 | chr07 | 4881911 | 4881919 | GCGTATATT | G | DEL (8) | upstream_gene_variant | | | 1\|1 | ✓ |
| 43 | IRB3517-3 | chr08 | 23894794 | 23894794 | A | G | SNV | 3_prime_UTR_variant | Os08g0483900 | Helix-loop-helix DNA-binding domain containing protein. | 0\|1 | |
| 44 | IRB3517-3 | chr09 | 7224224 | 7224224 | A | G | SNV | missense_variant | Os09g0297400 | [PPT1] Similar to Phosphate/phosphoenolpyruvate translocator. | 1\|1 | ✓ |
| 45 | IRB3517-3 | chr09 | 13007477 | 13007477 | G | A | SNV | synonymous_variant | | | 0\|1 | ✓ |
| 46 | IRB3517-3 | chr11 | 1188438 | 1188438 | C | T | SNV | 5_prime_UTR_variant | Os11g0126250 | Hypothetical gene. | 1\|1 | |
| 47 | IRB3517-3 | chr11 | 7365282 | 7365282 | T | C | SNV | missense_variant | Os11g0238000 | Similar to NB-ARC domain containing protein. | 1\|1 | ✓ |
| 48 | IRB3517-3 | chr12 | 3883429 | 3883435 | TCACTAC | T | DEL (6) | downstream_gene_variant | Os12g0176800 | Similar to Heterochromatin protein (Fragment). | 0\|1 | |
| 49 | IRB3790-2 | chr01 | 5274282 | 5274283 | GC | G | DEL (1) | intron_variant | | | 0\|1 | |
| 50 | IRB3790-2 | chr03 | 26252641 | 26252642 | A | AT | INS (1) | upstream_gene_variant | | | 0\|1 | |
| 51 | IRB3790-2 | chr04 | 12393088 | 12393088 | G | A | SNV | intergenic_region | | | 1\|1 | |
| 52 | IRB3790-2 | chr06 | 9338213 | 9872884 | TATG….TTGG | TACA….TCAT | INV (535k) | frameshift_variant [8] | Os06g0275000 [7] | Photoperiod-sensitivity-1, HEADING DATE 1 [7] | 0\|1 | ✓ |
| 53 | IRB3790-2 | chr07 | 23822057 | 23822057 | T | C | SNV | missense_variant | Os07g0586200 | [GELP92] Similar to Esterase precursor (EC 3.1.1.-) (Early nodule-specific protein homolog) (Latex allergen Hev b 13). | 0\|1 | |
| 54 | IRB3790-2 | chr08 | 21594849 | 21594853 | CTTAA | C | DEL (4) | upstream_gene_variant | | | 1\|1 | ✓ |
| 55 | IRB3790-2 | chr09 | 1369829 | 1369831 | GGA | G | DEL (2) | intergenic_region | | | 1\|1 | |
| 56 | IRB3790-2 | chr12 | 4637905 | 4637905 | G | A | SNV | upstream_gene_variant | | | 1\|1 | ✓ |

[1] Mutation No. is identical to that in Table 2.

[2] For the large deletions and the inversion, only first and last 4 bases are shown.

[3] Size of InDels is indicated in parentheses.

[4] Mutations putatively make high and moderate impact on protein function are indicated by orange and blue background color.

[5] "0|1" is heterzygous and "1|1" is homozygous.

[6] Mutations verified by Sanger sequencing were marked with checkmark. Primer sequences are shown in S1Table.

[7] Independent mutation calls are unified and counted as one RPL mutation because they are overlapped or located consecutively (Fig. 3).

[8] Evaluated manually.
